# Evolutionary Advantage of the Diversity-Generating Retroelement Hypermutating System

**DOI:** 10.64898/2026.02.18.706561

**Authors:** Léo Régnier, Francesco Amoroso, Paul Rochette, Raphaël Laurenceau, David Bikard, Simona Cocco, Rémi Monasson

## Abstract

Diversity-Generating Retroelements (DGRs) create rapid, targeted variation within specific genomic regions in phages and bacteria. They operate through stochastic retro-transcription of a template region (TR) into a variable region (VR), which typically encodes ligand-binding proteins. Despite their prevalence, the conditions under which maintaining such hypermutating system is favorable remain unclear. Here we introduce a two-timescale framework separating fast VR diversification from slow TR evolution, allowing the dynamics of DGR-controlled loci to be analytically understood. Combining data analysis and analytical calculations we quantity the fitness gain provided by the diversification mechanism of DGR with respect to standard mutagenesis. Our framework accounts for observed patterns of DGR activity in human-gut *Bacteroides* and clarifies when constitutive DGR activation is evolutionarily favored.

## Introduction

Diversity-Generating Retroelements (DGRs), first identified in the *Bordetella* phage [1], represent an important class of hypermutation systems that enable rapid adaptation in phage and microbial populations. Recent work has clarified their structural organization [2, 3], documented their diversity across microbial genomes [4–6], elucidated their biological roles [6, 7], and highlighted their potential for engineering applications [8, 9]. DGRs introduce mutations by copying a template region (TR) into a target variable region (VR) through an error-prone reverse transcription process [10–12] (Fig. 1**(a)**). The mechanism is inherently asymmetric. While the TR slowly evolves through standard mutagenesis, the VR is quickly and repeatedly diversified and overwritten, with mutations concentrated at positions corresponding to a specific TR nucleotide (typically, Adenine). This localized hypermutating system can generate extensive phenotypic diversity at loci involved in host recognition, cell–cell interactions, or environmental sensing [5–7].

**FIG. 1.**
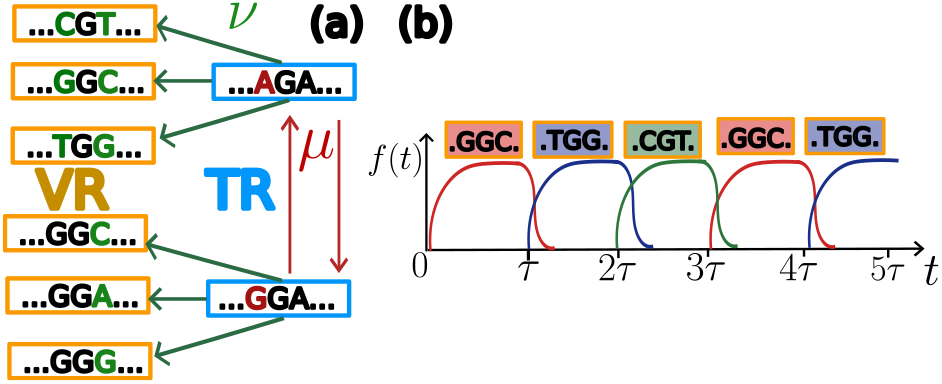
The DGR system. **(a)** The template region (TR, blue), after reverse transcrition with adenine-specific random mutations, replaces the variable region (VR, orange) at rate *ν*. Independently, VR and TR bases mutate at rate *µ*. **(b)** Sketch of a fluctuating environment. Every interval *τ*, the environment switches to favor a different target (VR) sequence. The population frequency *f* (*t*) of the fittest sequence (shown on top) then adapts to these shifts.

Recent studies on the human gut microbiome [5, 13, 14] illustrate the benefits of high DGR mutation rates [15]. Infants are born with very few DGRs in their microbiome, but by one year, they reach adult-like levels. Remarkably, ∼ 72% of the 388 parental DGRs were observed to switch their VR during this period. Some *Bacteroides* species (notably *B. ovatus* and *B. finegoldii* ) maintain high DGR activity: 13–40% of VRs diverge within 14 days. These results hint at the role of DGRs as accelerators of adaptive evolution, promoting the emergence of fitter variants on a dedicated timescale far faster than standard mutational processes.

Although hypermutation has been studied in other biological contexts [16–18], the conditions under which the distinctive architecture of the DGR hypermutating system is actually advantageous remain unclear. Furthermore, impairment (through mutations) of the reverse transcriptase could in principle silence diversification, highlighting a potential evolutionary fragility of DGRs. Hereafter, we propose an explicit mechanistic modeling to understand the long-term selective benefit of such constitutively active diversification.

### DGR model and evolutionary dynamics

Hereafter, a DGR is defined by the nucleotidic sequences of its template and variable regions, which we denote by, respectively, **VR** and **TR**; both sequences have the same length *L*. The **VR** is replaced through to the DGR mechanism at rate (probability per unit of time) *ν*, see Fig. 1**(a)**. In this process, nucleotides *C, G, T* in the **TR** are copied as such at the same positions in the **VR**, while *A*’s are equally likely to mutate into *C, G, T* or to remain unchanged [19] In addition, under point mutation process, each nucleotide in the **VR** and **TR** mutate with rate *µ*(≪ *ν*).

As proteins are expressed from the gene sequence including the **VR**, the fitness *S* (growth rate) of an individual—defined by its (**TR, VR**) pair—depends solely on the **VR**. We assume that *S* is additive across sites (no epistasis) [20, 21] as this simple model can capture a wide range of biological scenarios [22, 23]. We also assume that fitness contributions depend solely on the nucleotide identity at each position rather than the encoded amino acids. The fitness at time *t* is given by

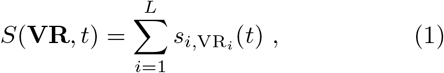

where *s*_*i,q*_ denotes the contribution of a nucleotide *q* = *A, T, C, G* at position *i* along the **VR** sequence. To account for possible environmental changes, we allow the *s* factors to explicitly depend on time *t*.

### Experimental background

The authors of [14] studied five Bacteroide DGRs, and found two species that maintained a high level of DGR activity both *in vitro* and *in vivo* in the absence of selective pressure (as evidenced by sustained activity in monocolonized mice).

After *t* =14 days, a fraction *φ* of **VR** ranging from 13 to 40 % (*in vitro*) and from 0.6% to 1% (*in vivo*) had diverged from the initial ones. Within our modeling framework, this fraction is simply given by *φ* = 1 − *e*^*−νt*^ due to the lack of selection (*s* independent of the nucleotide content), see details in Supplementary Materials (SM) [24] Sec. 1. Matching this formula with the experimentally observed fraction, we infer the DGR replacement rates for these two Bacteroides to range between *ν* = 10^*−*2^ day^*−*1^ and 4. 10^*−*2^ day^*−*1^ from *in vitro* data, and between 7. 10^*−*4^ day^*−*1^ and 2. 10^*−*3^ day^*−*1^ from *in vivo* data, see Tab. I.

As a point of comparison, the actual doubling time of Bacteroides is around 2 hours in optimal conditions and up to one day [25, 26] (when selection plays a role), resulting in a growth rate *S* of the order of 1 to 10 day^*−*1^.

Equating the length of the DGR sequences with the number of variables sites, *i*.*e*. of adenines in the **TR**, we estimate *L* ≃ 13 − 46 from Ref. [5] (confidence interval with 95% of the data, see SM Sec. 2). As the spontaneous base mutation rate *µ* is of the order of 10^*−*9^ − 10^*−*8^ day^*−*1^ [26, 27], we obtain that the rate *µL* of spontaneous mutations in the **TR** (and in the **VR** in the absence of DGR) are several orders of magnitude smaller than the DGR replacement rate *ν*, see Tab. I.

### VR selection at fixed TR

On time scales *<* 1*/*(*µL*), no mutation happens on the **TR**, while the **VR** evolves under selection and is periodically overwritten via DGR-mediated diversification at rate *ν*. We hereafter study analytically this fixed **TR** regime, in the case of a time-dependent environment. For the gut microbiota, the intestinal microbial community changes over a time scale ranging from days (e.g., in response to dietary changes) to years, as documented in previous studies [28–31]. Environmental changes can also result from the presence of other DGR populations — either constitutionally diverse (as in *Bordetella*) or arising from other bacterial species.

For the sake of mathematical simplicity, we consider the environment to be periodic [32–34]; note that the main results, such as the asymptotic population growth rate, still hold when relaxing part of this hypothesis, *e*.*g*. when randomly drawing the next environment, see SM Sec. 4.C. Let us call 4*τ* the period of the environmental changes (Fig. 1**(b)**). For *t* ∈ [*q*^*′*^*τ*, (*q*^*′*^ + 1)*τ* ] with *q*^*′*^ = 1, …, 4, the preferred nucleotide at position *i* is *q*^*∗*^(*t*) = *q*^*′*^ (such that two successive environment do not select for overlapping sequences) and confers a fitness advantage *s >* 0 with respect to the other three ones, *i*.*e*. 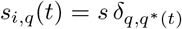 in Eq. (1). The population sizes *n*_**VR**_(*t*) of **VR** at time *t* obey the Wright-Fisher equations (finite-size effects are neglected, due to large populations of Bacteroides species, around 10^11^ − 10^14^ per gram in the gut microbiome [35])

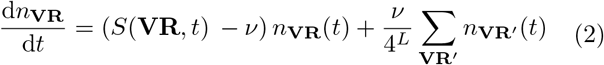

Resolution of this set of linear differential equations is detailled in SM Sec. 4. After each period, the 4^*L*^- dimensional vector of the population sizes *n*_**VR**_ is multiplied by a (4^*L*^ × 4^*L*^)-dimensional evolution matrix, whose entries encode the transition rates appearing in Eq. (2). The effective growth rate then reads *S*_*L*_ = ln Λ*/*(4*τ* ), where Λ is the largest eigenvalue of the evolution matrix. We find that it is approximately given by

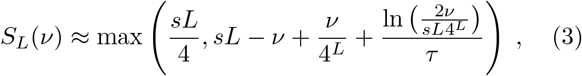

see SM Sec. 4 for the full derivation.

Results are shown in Fig. 2**(a)**, and are in agreement with large-size population dynamics simulations. The growth rate is maximal when the rate *ν* is roughly equal to 1*/τ* [32, 33]. When the DGR substitution rate *ν* is lowered, *S*_*L*_ decays (slowly for large *τ* ): the system has less time to adapt to the fluctuating environment as mutations become rarer. Conversely, *ν* should not exceed *s*, as overly frequent DGR-generated replacements would reduce the adaptive benefit. This is illustrated by Fig.1**(b)**: large *ν* (fast adaptation) leads to rapid growth of the optimal sequence frequency *f* (*t*) but to a low value once it stabilizes.. Notice that *τ* must not be significantly larger than 1*/s*, as a low spontaneous mutation rate *µ* would suffice to track slow environmental changes. These conditions align with experimental observations [14], where *ν* ≲ *s*. For *L* of the order of 20 (the effective number of variable sites) and *s* = *S*_*e*_*/L*, this ensures both sufficient positive selection and high growth rates (Table I).

**FIG. 2.**
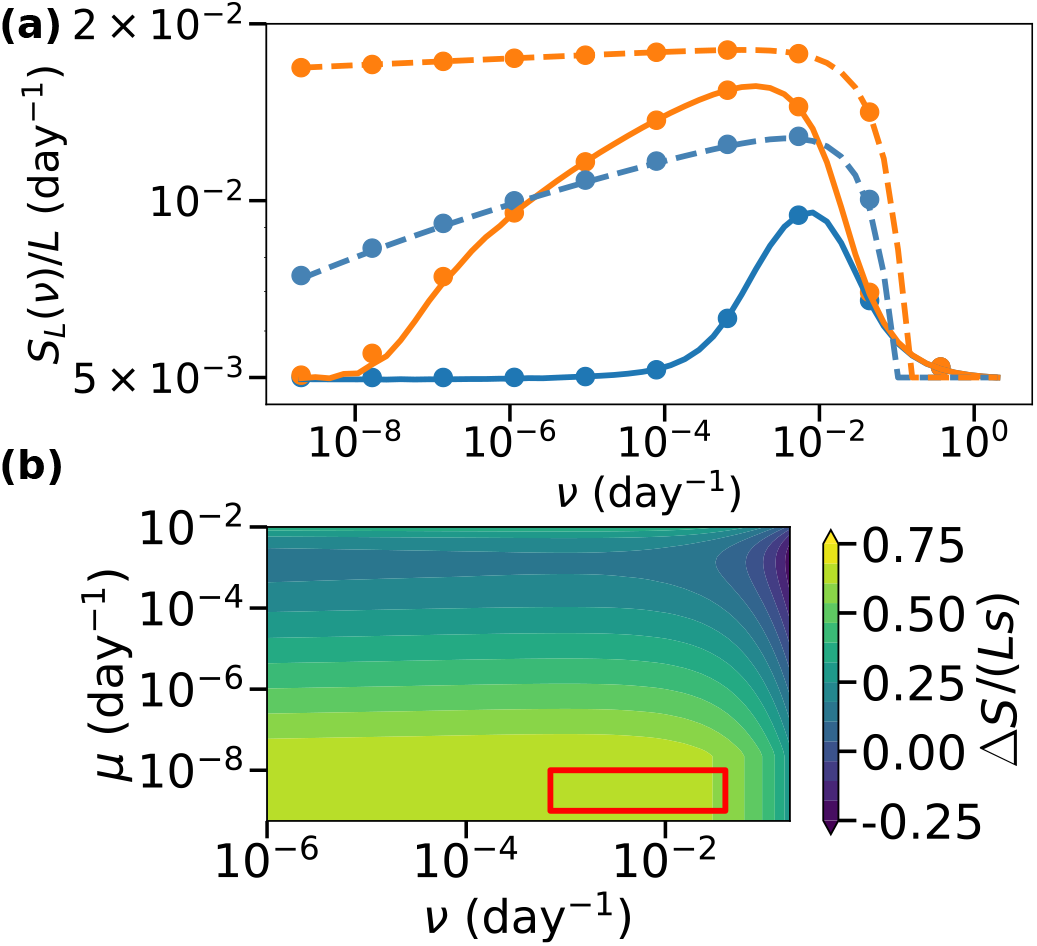
VR selection. **(a)** Effective growth rate vs. DGR replacement rate *ν*. Plain lines: *L* = 1 (population dynamics), dashed lines: *L* = 10 (approximation in Eq. 3), points: exact solution of Eq. 2. Blue: *τ* = 250, orange: *τ* = 1000. **(b)** Phase diagram of DGR prevalence in the (*ν, µ*) plane. Color shows the fitness difference Δ*S* (normalized by *Ls*) between DGR and non-DGR systems for *τ* = 10^3^, *L* = 22. Parameter values: *s* = 2 *×* 10^*−*2^, population size *N* = 10^8 a^. The red box delimits the range of *ν* and *µ* given in Tab. I. Time unit is day, rates are in day^*−*1^. ^a^ *Nμ* and *Nν* much larger than the inverse discretization time step for most of the values considered.

**TABLE 1.** Rates for DGR dynamics in *Bacteroides*.

| Quantity | Notation | Range ($\text{day}^{-1}$) |
| --- | --- | --- |
| DGR replacement rate | $\nu$ | $7 \cdot 10^{-4} - 4 \cdot 10^{-2}$ |
| Growth rate | $S$ | $1 - 10$ |
| Growth rate per variable site | $s$ | $2 \cdot 10^{-2} - 1$ |
| Spontaneous mutation rate | $\mu L$ | $10^{-8} - 5 \cdot 10^{-7}$ |

In Fig. 2**(b)**, we map the parameter space where DGR-equipped populations outcompete non-DGR populations, *i*.*e*. where the fitness difference Δ*S* = *S*_*L*_(*ν*) − *LS*_1_(*µ*) is positive. For the range of *ν* and *µ* values given in Tab. I (red box in Fig. 2), we find Δ*S >* 0. Equivalently, Δ*S* can be interpreted as the maximal metabolic or functional burden –in terms of deficit in growth rate– the DGR mechanism can afford to remain competitive with respect to standard mutagenesis. This DGR-associated burden could be experimentally estimated through the difference of growth rates in a competition assay including a population with DGR and another without DGR (after inactivation of the RT [14]) in the absence of selection on the VR.

### Comparison with experimental data

We next compare our theory to the *in vitro* gut microbiota dataset of a *Bacteroides ovatus* population from [14].

In this experiment, a mouse was inoculated with a DGR population containing a single **VR** (the parental sequence) and an *Altered Schaedler Flora* (ASF, a small bacteria consortium with competition) microbiome to enable selection; mutations over *L* = 22 TR adenine positions were detected in the **VR** population. Accounting for the emergence of new sequences via random replacements and binomial sampling [23], we estimate a fitness model *S*(**VR**), where each nucleotide contributes independently to the growth rate of the sequence, see SM Sec. 3 for details. We then compare the experimental fitness of non-parental **VR**s to numerical simulations of: *(i)* A DGR population with a DGR replacement rate *ν* = 10^*−*3^, and *(ii)* Populations with a per-base mutation mechanism at rates *µL* = 10^*−*3^ (equal to the rate *ν* of new VR sequence appearance) and *µ* = 10^*−*8^ (expected rate for standard mutations). As observed in Fig. 3, the DGR population outcompetes the non-DGR population even with similar mutation rates. Furthermore, the numerical simulations are in excellent agreement with the experimental curve, confirming the validity of the independent–site model for both DGR diversification and fitness in this time range.

**FIG. 3.**
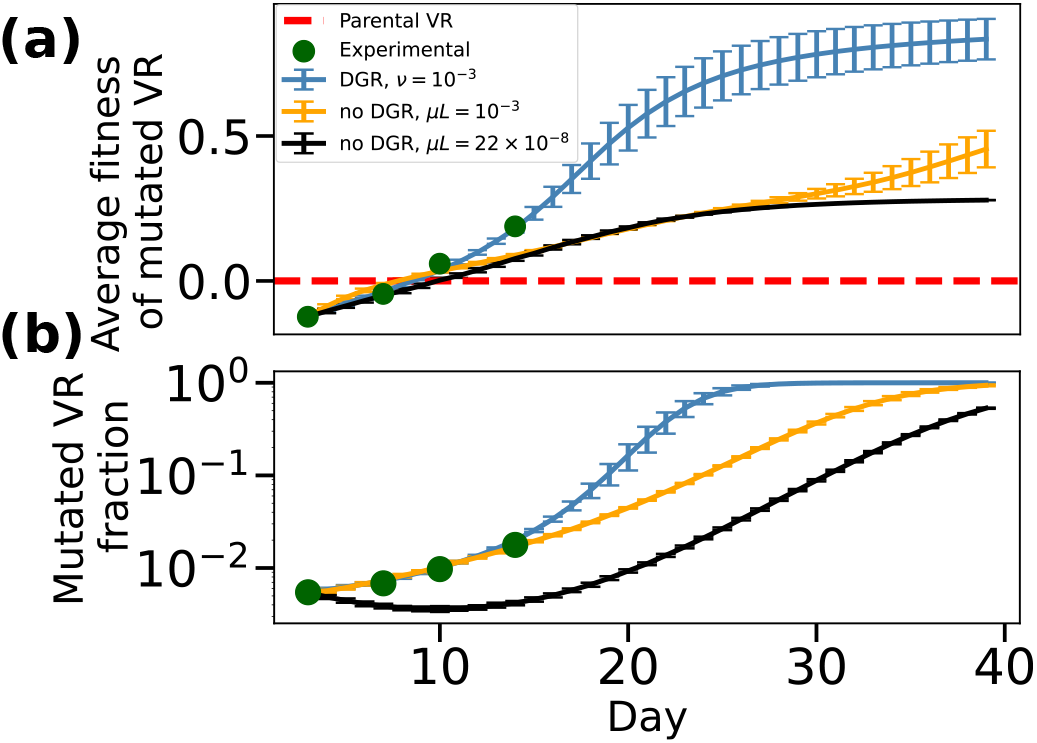
Population dynamics of *Bacteroides ovatus* in the gut. We compare **(a)** the average fitness of mutants (with variable regions differing from the initial strain) and **(b)** their observed frequency in the mouse gut microbiota in the presence of ASF across three mutation scenarios: DGR replacement mechanism at rate *ν* = 10^*−*3^ (blue), per-base mutation mechanism at the same rate *µL* = *ν* (orange), and per-base mutation mechanism at rate *µ* = 10^*−*8^ (black). Experimental data points are shown as green dots. The parental VR fitness is represented by a red dashed line. Error bars show standard deviations over 100 simulations.

### TR selection through mutations

We now allow the nucleotides in the **TR** to mutate at a (low) rate *µ*. A detailed treatment of the Wright-Fisher equations describing the evolution of the population of (**TR, VR**) pairs in a fluctuating environment is provided in SM Sec. 6. We summarize the main results below.

A **TR** sequence can be evolutionarily fit in distinct ways. First, it may carry many *A*’s that allow the **VR** to diversify and adapt to any environment. However, continual diversification of the **VR** at a rate *ν* leads to the frequent loss of adaptation in stable contexts. Second, the **TR** may carry non-Adenine nucleotides that are well-adapted to the current environment; their copies in the **VR** are far less mutable (rate *µ* ≪ *ν*) and thus provide a fitness advantage if the time scale for environmental switch is long. Third, purely random sequences may be favored for entropic reasons through accumulation of mutations when *µ* is large compared to 1*/L*. The phase diagram showing the prevalence of the three regimes in the (*L, τ* ) plane is given in Fig. 4**(a)**. The boundaries of the phases can be informally retrieved as follows. The maximally-diversifying **TR** (with all adenines) competes with the best adapted sequence in the current environment, whose population grows as 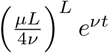 : the prefactor corresponds to the *L* spontaneous mutation events needed to reach the best adapted **TR**, and the exponential term the gain in losing the DGR (SM Sec. 6). Notice that if the environment favors *A* on the **VR**, then this adapted nucleotide can be produced only by a *A* on the **TR**. This case does therefore not contribute to the loss of Adenines on the **TR** (SM Sec. 6). In addition, random sequences invade the population, with a fraction growing as *e*^*µLt*^. As a consequence, the all-*A* TR is favored for quickly-fluctuating environments, and outcompeted when the switching time *τ* exceeds

**FIG. 4.**
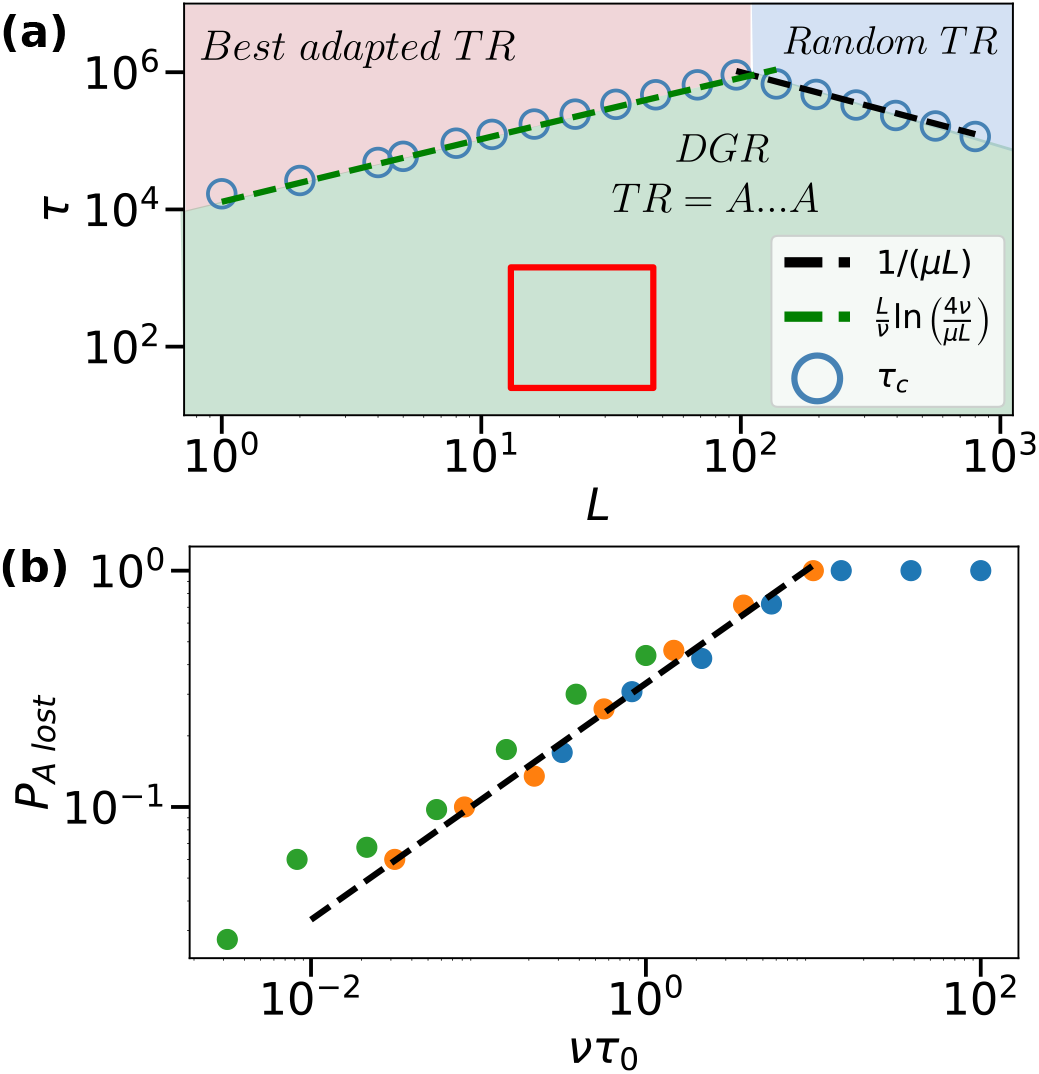
TR selection. **(a)** Phase diagram in the (*L, τ* ) plane. Dashed lines show the theoretical prediction for *τ*_*c*_ in Eq. (4). Blue points are determined by solving the dynamics for infinite population size. Parameters are *µ* = 10^*−*8^ and *ν* = 10^*−*3^. The red square delimits the region *τ ∈* [1*/*(4 *×* 10^*−*2^); 1*/*(7 *×* 10^*−*4^)] and *L ∈* [13; 46] (see Tab. I). **(b)** Probability *P*_*A lost*_ that the fraction of **TR** with an *A* (initially, equal to 1) is lower than 1*/*4 as a function of *ντ*_0_ where *τ*_0_ is the typical duration of an environment. Dashed line corresponds to (*ντ*_0_)^*β*^ . We take *µ* = 10^*−*4^, *β* = 1*/*2 and *ν* = 10^*−*3^ (green), 10^*−*2^ (orange) and 10^*−*1^ (blue).

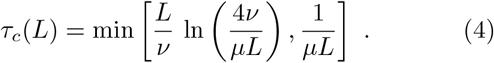

see Fig. 4**(a)**. An important consequence of this result is that the DGR phase is best stable over time for the optimal length (number of A’s) given by

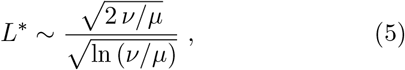

Using the in vivo estimate of *ν* with the estimate of *µ* given in Table I, we obtain *L*^*∗*^ spanning values of the order of 10^2^ − 10^3^. This result stresses that having extended variable regions in the **TR** can be beneficial for the resilience of the DGR system, but that too long **TR** are not robust against spontaneous mutations. This is consistent with the relatively low number *L* (up to 10^2^) of variable sites found in [5, 14] and displayed in SM Sec. 2. We stress that our analysis focuses on the potential loss of DGR due to the TR dynamics only. In practice, *L*^*∗*^ could be further limited by other factors, such as reverse transcriptase inactivation, a cost of DGR activity scaling with *L*, or structural constraints of the residues to diversify, see **Conclusion and extensions**.

The results above may break down if the environmental changes are not close to being periodic, *i*.*e*. if the distribution of switching times *τ* has fat tails extending well above 1*/ν*. To quantitatively understand this phenomenon we focus on the simplest case of TR sequences of length *L* = 1. The overwhelming majority of **TR** sequences either carry a *A*, or the nucleotide best adapted to the current environment, in agreement with Fig. 4**(a)**. The fraction of the latter is approximately given by

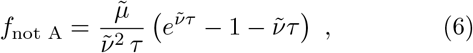

where 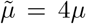 and 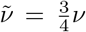 We compare this approximation with simulations in SM Sec. 6, confirming its validity. Adenines in the **TR** become strongly disfavored if *τ* may significantly exceed 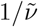 according to Eq. (6). For example, consider the case of *τ* algebraically distributed, with typical time-scale *τ*_0_ and exponent *β, i*.*e*. with distribution ℙ (*τ* ≥ *t*) = (*τ*_0_*/t*)^*β*^. The adenine is lost with a probability of the order of 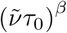, condition which corresponds to *f*_not A_ ∼ 1. This theoretical scaling is confirmed in Fig. 4**(b)** by simulations estimating the probability that, by the end of the environmental bout of random duration *τ*, the population of **TR** carrying *A* no longer dominates (fraction smaller than 1*/*2). This example shows that strongly (algebraically) variable environmental duration can be disastrous for the maintenance of the DGR mechanism, despite the fact that the typical timescale *τ*_0_ of these environmental duration still obey 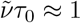.

### Conclusion and extensions

Our analysis offers a framework to understand the conditions under which the DGR system offers fitness advantages over standard spontaneous mutation processes in fluctuating environments. If the environmental context is stable or does not vary quickly enough, the DGR becomes progressively inoperant through the loss of the Adenines in the **TR** after some critical time we have estimated.It is thus crucial to identify the sources of environmental fluctuations, whether they are extrinsic (e.g., temperature shifts, changes in food availability) or intrinsic (e.g., regulation by other populations, with or without DGRs).

An illustration of the latter is provided by two competing DGR populations. Preliminary investigation shows that an effective time scale *τ* for environmental changes emerges, as a complex function of time delays (e.g., spatial distributions, reaction times of one population to changes in the other, or biological processes) [36–39], as well as their respective DGR replacement rates, see SM Sec. 5.B for details.

One promising experimental direction would be to extend the study of Ref. [14] to observe whether, over the long term, DGR-equipped populations are outcompeted by non-DGR populations in the mouse ASF microbiome, or the dominant VR changes periodically indicating environment fluctuations even in this simplified system. Alternatively, DGR activity could be lost due to random mutations in the reverse transcriptase gene, a scenario that warrants further modeling. Scenarios in which finite population size modulates the benefits of hypermutation could also be considered [17].

To prevent the emergence of low-fitness mutants, organisms could also tune DGR activation and deactivation despite costs induced by sensing environmental fluctuations [14, 40]. Regulation could be achieved by downregulating DGR when unnecessary [5, 14], silencing the system via mutations impairing reverse transcriptase [11, 14, 41], or in engineered systems, bypassing the VR entirely to reintegrate cDNA directly into the TR [8]. Inspired by persister cell mechanisms [42, 43] and recent observations [14], we refine our model by introducing activation and deactivation rates for DGR activity, see SM Sec. 5.A. Our analysis reveals that populations can oscillate between active and inactive DGR states in response to environmental changes, often remaining inactive for extended periods. This may explain the rarity of highly active DGR systems in the three inactive strands of Ref. [14], as such variants may only be transiently selected. We predict that experiments with frequent environmental fluctuations or conditions counter-selecting the parental VR sequence would increase their detectability.

Future analysis should take into account the existence of mutational biases inherent to DGRs and their consequences for protein-level selection. Because selection acts on amino acids, the position of adenines within codons can substantially affect the number of accessible amino-acid variants (for example, AAC codons can diversify into 15 amino acids, whereas having an adenine only at the third position yields negligible variation), leading to an effectif epistatic effect at the nucleotide level. These positional effects, together with biases in DGR A-to-N substitutions [8, 9, 14], shape the landscape of possible receptor or ligand variants. Remarkably, the presence of epistatic effects may not affect the dynamics of exploration of DGR. We report in SM Sec. 5.C preliminary results about the exploration of a simple epistatic landscape that depend solely on the distance to the optimal VR. Our analysis shows that DGR population growth remains unaffected by the landscape shape, whereas per-base mutation mechanisms are strongly impacted as local evolutionary dynamics can be trapped in local fitness maxima, *e*.*g*. in the presence of sign epistasis. This result hints at the superiority of DGR populations to explore rugged fitness landscapes with respect to standard mutagenesis.

## Data availability

The code for simulations and figures is available in the GitHub repository: https://github.com/LeoReg/DGR_pop_dynamics.git.

## Funding informations

This work was supported by the French National Research Agency (ANR) project Prodigen under Grant No. ANR-23-CE45-0034.

