## Supplementary Material for "Evolutionary Advantage of the Diversity-Generating Retroelement Hypermutating System"

### SUPPLEMENTAL MATERIAL
### A model of Diversity Generating Retroelements population dynamics.

CONTENTS


S1. Experimental background 1

S2. Estimation of the number $L$ of Adenines going through DGR diversification 2

S3. Analysis of the mice dataset 3
- A. Details of the procedure 3
  - 1. Inference of parameters from the experimental data 3
  - 2. Simulation of the population dynamics 4
  - 3. Estimation of the average fitness of mutants 6

S4. Fast dynamics of the VR population at fixed TR 6
- A. $L = 1$ 6
- B. $L > 1$ 7
  - 1. $Q = 2$: perturbation theory 7
  - 2. General $Q$ lower bound 11
- C. Stochastic environment 12

S5. Model extensions 15
- A. On-off DGR 15
- B. Competing populations 15
  - 1. Two populations, two genotypes 16
  - 2. Extended genotypes case 17
- C. Epistatic landscape: single genotype selection 19
  - 1. Single genotype selected 19
  - 2. Generalization 20

S6. Slow dynamics of the TR 21
- A. $L = 1$ 21
  - 1. The **TR**=A population: first order 21
  - 2. The **TR**$\neq$A population: the order $\mu$ correction 23
  - 3. Simplified dynamics 27
  - 4. Dicussion about **VR**=A selection by the environment 27
- B. $L > 1$ 28

Supplementary References 31


#### S1. EXPERIMENTAL BACKGROUND

In this section, we explain the calculations of the **Experimental background** section of the main text.
The setup is the following: the experiment starts with a population $n_0(0)$ of individuals with the same Variable Region (VR). At a rate $\nu$, the Template Region (TR) go through the Diversity Generating Retroelement (DGR) mechanism which leads to the replacement of the initial VR by a new one. We note the population in this category $n_1(t)$, population initially empty.
Because there is no selection against the VR in the experiment of Ref. [S1], we suppose that both populations grow at a same rate $S_e$. Finally, because it is unlikely for the system to produce the same VR as the initial one through the DGR mechanism, the number of possible VR growing exponentially with the number of Adenines, we consider that an individual in population 1 stays there. Of note, even for small number of Adenines, this changes the estimated rate

$\nu$ by a prefactor $1 - 1/Q$ at most (probability to produce the same nucleotide with only one diversifying nucleotide). From this, we have the differential equations describing the time evolution of the system:

$$\dot{n}_0(t) = (S_e - \nu)n_0(t) \tag{S1}$$

$$\dot{n}_1(t) = S_e n_1(t) + \nu n_0(t) \; . \tag{S2}$$

The solution of this system of equations is given by:

$$n_0(t) = n_0(0)e^{(S_e-\nu)t} \; , n_1(t) = n_0(0)\left(e^{S_e t} - e^{(S_e-\nu)t}\right) \; , \tag{S3}$$

from which we deduce the expression of the fraction $\varphi$ of the population with distinct **VR** from the original one as a function of time given in the main text:

$$\varphi = \frac{n_1(t)}{n_0(t) + n_1(t)} = 1 - e^{-\nu t} \; . \tag{S4}$$

We analyze the data from Ref. [S1] to get the fraction of DGR generated sequences (sequences with mutations at adenine locations in the TR). A strong filtering procedure is applied in order to remove all sequences containing sequencing errors (namely 'N' nucleotides) and sparse mutations, constant within the dataset, not attributable to the DGR generation process. The estimation of $\nu$ for the in-vivo cases is performed using data collected from stool samples of a mouse monocolonized with B. Ovatus (sample details are reported in Supplementary Table 1) and is described more precisely in S3 A 1.

#### S2. ESTIMATION OF THE NUMBER $L$ OF ADENINES GOING THROUGH DGR DIVERSIFICATION

We do an estimate of the number of variable sites in the region undergoing DGR via the dataset assembled in Ref. [S2]. Because it is not the whole TR that will go through the hypermutation mechanism of the DGR, we consider only nucleotides which are between two mutations (detected in the VR). All Adenines are then considered to be potential variable sites (even though there was no mutation in the VR at those positions). The histogram of the Adenine count $L$ is presented in Fig. S1. We obtain that 95% of sequences have less than $L = 41$ Adenines in the region undergoing the DGR hypermutation mechanism. On top of that, we also display the distribution of the number of nucleotides (or VR length) in the region undergoing DGR, for which more than 95% of sequences are of length smaller than 126.

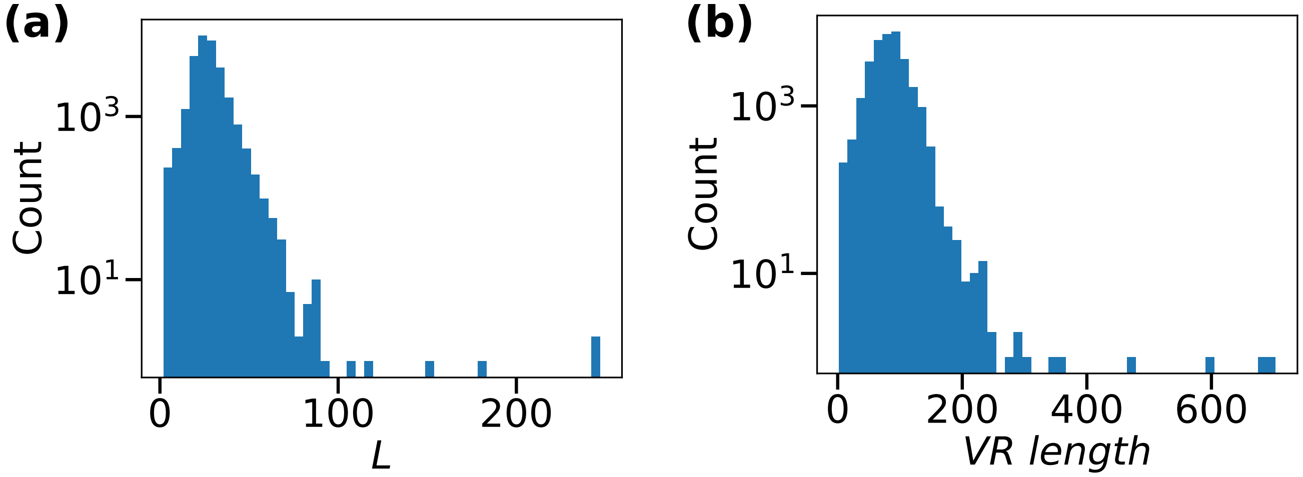


Fig. S1. **Distribution of Adenines and nucleotides count in the diversified region.** Histogram of **(a)** the number $L$ of Adenine **(b)** the number of nucleotides (or VR length) in the region detected as DGR from Ref. [S2], with the additional constraint that the nucleotides are in an interval where DGR mutations have been detected

#### S3. ANALYSIS OF THE MICE DATASET

##### A. Details of the procedure

###### 1. Inference of parameters from the experimental data

By considering the dataset of Ref. [S1], we want to model the VR population dynamics of the Bacteroides Ovatus species. To do so, we infer:

- the VR generation probability, denoted in the following by $\pi_{DGR}(\boldsymbol{VR})$, from the in-vitro data, where no selection occurs. We infer it using an independent site model of parameters $(g_{i,q})_{1\leq i\leq L, q\in\{A,T,C,G\}}$,

  The inference dataset is retrieved by merging the sequences mutated by DGR and observed at day 3 in two different bacterial cultures showing similar mutational profiles. The cultures are grown in two different tubes (see [S1] for the detailed procedure); the specific samples used are listed in Supplementary Table 1.The merging results in a total of $\sim$ 3000 sequences and 22 adenine positions along the TR parental that show sufficient diversification. The components of $\boldsymbol{g}$ are computed as follows:

  $$g_{i,q} = log(f_{i,q} + \epsilon)$$

  where $f_{i,q}$ is the observed frequency of nucleotide $q$ at position $i$, and the pseudocount $\epsilon = \frac{1}{N_{vitro}}$, used to avoid zero-frequencies cases. $N_{vitro}$ is the total number of in vitro non-parental VR sequences used for the inference. Up to a normalization factor, $\pi_{DGR}(\boldsymbol{VR})$ is given by:

  $$\pi_{DGR}(\boldsymbol{VR}) \propto \exp\left(\sum_{i=1}^{L} g_{i,VR_i}\right) \tag{S5}$$

  .

- the DGR VR replacement rate $\nu$ in-vivo in the monocolonized mouse (without the ASF, and thus without induced selection, nor any significant natural selective pressure). In the following, we take $\nu = 10^{-3}\ day^{-1}$, because this value is representative of an intermediate $\nu$ among the ones inferred in vivo at the different experimental timepoints (see Supplementary Table 1 for the list of samples used):

  - $2.3 \times 10^{-3}\ day^{-1}$, inferred at day 3
  - $1.1 \times 10^{-3}\ day^{-1}$, inferred at day 7
  - $9.4 \times 10^{-4}\ day^{-1}$, inferred at day 10
  - $6.5 \times 10^{-4}\ day^{-1}$, inferred at day 14

- the fitness landscape $S(\boldsymbol{VR})$, where we assume that each site contributes independently, so that

  $$S(\boldsymbol{VR}) = \sum_{i=1}^{L} s_{i,VR_i}\ , \tag{S6}$$

  , from the in-vivo time series dataset with ASF, where a strong selective pressure is observed (see Supplementary Table 1 for the detailed list of the datasets used for the inference). Similarly to Ref. [S3], the fitness landscape is inferred by maximizing the following log-likelihood:

  $$\mathcal{L} = \sum_{i=0}^{T-1} \mathcal{L}^t(t_{i+1} - t_i) \qquad \mathcal{L}^t(\Delta t) = \log\left[P\left(\boldsymbol{n}(t+\Delta t) \mid \boldsymbol{n}(t)\right)\right] \tag{S7}$$

  where $\{t_1, ..., t_T\}$ is the sequence of experimental time points (in days), $\mathcal{L}^t(\Delta t)$ is the likelihood at fixed time point $t$, associated with the time interval $\Delta t = t_{i+1} - t_i$, and $\boldsymbol{n}(t)$ is the population vector at time $t$, with $n_{\boldsymbol{VR}}(t)$ being the number of sequences $\boldsymbol{VR}$ observed in the population at time $t$. We note $\mathcal{S}_t$ the set of $\boldsymbol{VR}$ present at time $t$ in the sample.

In this framework, $P(\boldsymbol{n}(t+\Delta t) \mid \boldsymbol{n}(t))$ represents the probability of observing the population $\boldsymbol{n}(t+\Delta t)$ at time $t+\Delta t$, given the population $\boldsymbol{n}(t)$ at time $t$ and is modeled as follows (up to a proportionality factor independent of the parameters $(s_{i,q})$ to infer):

$$P(\boldsymbol{n}(t+\Delta t) \mid \boldsymbol{n}(t)) \propto \prod_{\boldsymbol{VR} \in \mathcal{S}_{t+\Delta t}} p_s(\boldsymbol{VR}, \Delta t)^{n_{\boldsymbol{VR}}(t+1)} \tag{S8}$$

$p_s(\boldsymbol{VR}, \Delta t)$ is the so called *selectivity* of sequence $\boldsymbol{VR}$, and combines the DGR generation mechanism and the fitness growth in the following formula:

$$p_s(\boldsymbol{VR}, \Delta t) = \frac{\left[(1 - p_{DGR}(\Delta t))\, n_{\boldsymbol{VR}}(t) + \delta_{n_{\boldsymbol{VR}}(t),0}(1 - \delta_{n_{\boldsymbol{VR}}(t+\Delta t),0})\right] e^{S(\boldsymbol{VR})\Delta t}}{\sum_{\boldsymbol{VR}' \in \mathcal{S}_t \cup \mathcal{S}_{t+\Delta t}} \left[(1 - p_{DGR}(\Delta t))\, n_{\boldsymbol{VR}'}(t) + \delta_{n_{\boldsymbol{VR}'}(t),0}(1 - \delta_{n_{\boldsymbol{VR}'}(t+\Delta t),0})\right] e^{S(\boldsymbol{VR}')\Delta t}} \tag{S9}$$

where $p_{DGR}(\Delta t) = 1 - e^{-\nu \Delta t}$ is the fraction of DGR VR replaced sequences ($\nu = 10^{-3}$) during the time interval of duration $\Delta t$, and $S(\boldsymbol{VR})$ is computed according to (S6). The second term of the numerator accounts for the cases in which a new $\boldsymbol{VR}$ sequence appears in the population during the time interval $\Delta t$, due to the DGR generation process.

Due to the very small number of VR sequences, we limit ourselves to the additive site model for both DGR VR replacement and the fitness model, which is a good approximation in this limit [S4].

Supplementary Table 1. Details of the samples used for the inference.

| **Sample** | **Experiment** | **SRA Accession** | **Day** | **Application** |
|---|---|---|---|---|
| Tube 1 (in vitro) | Exp. 15 | SRR31376262 | D3 | Inference of $\pi_{DGR}(\boldsymbol{VR})$ |
| Tube 3 (in vitro) | Exp. 15 | SRR31376260 | D3 | Inference of $\pi_{DGR}(\boldsymbol{VR})$ |
| Mouse 1, Cage 1 (in vivo, no ASF) | Exp. 6 | SRR31376296 | D3 | Inference of $\nu$ |
| | | SRR31376292 | D7 | Inference of $\nu$ |
| | | SSRR31376279 | D10 | Inference of $\nu$ |
| | | SRR31376274 | D14 | Inference of $\nu$ |
| Mouse 1, Cage 3 (in vivo, with ASF) | Exp. 15 | SRR31376287 | D3 | Inference of $S(\boldsymbol{VR})$ |
| | | SRR31376283 | D7 | Inference of $S(\boldsymbol{VR})$ |
| | | SRR31376239 | D10 | Inference of $S(\boldsymbol{VR})$ |
| | | SRR31376233 | D14 | Inference of $S(\boldsymbol{VR})$ |

##### 2. *Simulation of the population dynamics*

The simulation of the population dynamics takes into account three different scenarios, in order to highlight the role of the DGR mechanism in the average fitness of the mutant population. Hereafter we will use the same notation of the previous subsection, and by 'target positions', we will refer to the 22 adenine positions of S3 A 1

- Active DGR population: at each time step, starting from $t_i$, the population $\boldsymbol{n}(t_i)$ undergoes three consecutive events to reach $\boldsymbol{n}(t_{i+1})$:
  1. DGR generation process: given the generation rate $\nu = 10^{-3}$ and a time interval $\Delta t = t_{i+1} - t_i$, in each population $n_{\boldsymbol{VR}}(t_i)$, $\forall \boldsymbol{VR} \in \mathcal{S}_{t_i}$, $k$ sequences get replaced by new DGR-generated sequences. $k_{\boldsymbol{VR}}$ is modeled as follows:

$$k_{\boldsymbol{VR}} \sim \mathrm{Binom}(n_{\boldsymbol{VR}}(t_i), p_{DGR}(\Delta t)) \;, \tag{S10}$$

and the DGR generation process uses the $\boldsymbol{g}$ vector components to insert at each TR adenine position $i$ the nucleotide $q \in \{A, C, G, T\}$, according to the probability vector $\boldsymbol{p}_i$, with components

$$p_{i,q} = \frac{e^{g_{i,q} - \max(\boldsymbol{g}_i)}}{\sum_{q'} e^{g_{i,q'} - \max(\boldsymbol{g}_i)}} \ . \tag{S11}$$

The original population of $\boldsymbol{VR}$ gets updated as: $n_{\boldsymbol{VR}}(t_i) = n_{\boldsymbol{VR}}(t_i) - k_{\boldsymbol{VR}}$ and each one of the $k_{\boldsymbol{VR}}$ DGR generation processes leads to a new VR sequence, $\boldsymbol{VR'}$, that enriches $\mathcal{S}_{t_i}$ (the enlarged set can be denoted as $\mathcal{S}_{t_i,DGR}$), and contributes with a count of 1 to its population, $n_{\boldsymbol{VR'}}(t_i) = n_{\boldsymbol{VR'}}(t_i) + 1$.

2. Fitness-based growth: the population of each $\boldsymbol{VR}$ present in $\mathcal{S}_{t_i,DGR}$, grows according to the formula:

$$n_{\boldsymbol{VR}}(t_i) \leftarrow n_{\boldsymbol{VR}}(t_i) e^{S(\boldsymbol{VR})\Delta t} \ , \tag{S12}$$

where $S(\boldsymbol{VR})$ is computed according to (S6), using the inferred fitness landscape.

3. Multinomial sampling: in order to recover a final population of size $N$ that is constant for all the time points, we compute the normalized weights for the whole VR population as:

$$f_{\boldsymbol{VR}}(t_i) = \frac{n_{\boldsymbol{VR}}(t_i)}{n(t_i)} \ , \tag{S13}$$

with $n(t_i)$ being the total population size at time $t_i$ after the fitness-based growth. A multinomial sampling is then performed using the vector of normalized weights of the population at time $t_i$. The counts of the $\boldsymbol{VRs}$ in the population at time $t_{i+1}$ are sampled as:

$$(n_{\boldsymbol{VR}^{(1)}}(t_{i+1}), \dots, n_{\boldsymbol{VR}^{(n)}}(t_{i+1})) \sim \text{Multinomial}(N; f_{\boldsymbol{VR}^{(1)}}(t_i), \dots, f_{\boldsymbol{VR}^{(n)}}(t_i)) \tag{S14}$$

$$S_{t_{i+1}} = \{\boldsymbol{VR}^{(1)}, \dots, \boldsymbol{VR}^{(n)}\} \tag{S15}$$

- no DGR (spontaneous per-base mutation at rate $\mu$): when there is no DGR, mutant VRs can only be generated by a spontaneous mutation mechanism. We consider two values for $\mu$.
  In the first case, we consider the rate of spontaneous mutation per base $\mu$ for Bacteroides of the order of $10^{-9} - 10^{-8}\ day^{-1}$.
  In the second case, in order to look at the DGR generation mechanism and the spontaneous mutation process on comparable rate-scales, we force $\mu L = \nu = 10^{-3}$, so that, given $\Delta t$, $p_{DGR}(\Delta t) = p_{RM}(\Delta t)$.
  Again, starting from any population at time $t_i$, three consecutive events lead to the new population at time $t_{i+1}$:

1. Generation of spontaneous mutations: the formalism is analogous to the one of point S3 A 2, except for the following differences: in the absence of DGR, $p_{DGR}(\Delta t)$ becomes $p_{RM}(\Delta t) = 1 - e^{-(\mu L)\Delta t}$ ($RM$ stands for random mutation), which represents the probability of a given $\boldsymbol{VR}$ being randomly mutated; each $\boldsymbol{VR}$ population, of size $n_{\boldsymbol{VR}}(t_i)$, loses $k_{\boldsymbol{VR}}$ elements due to random mutation events, where $k_{\boldsymbol{VR}}$ is drawn as follows:

$$k_{\boldsymbol{VR}} \sim \text{Binom}(n_{\boldsymbol{VR}}(t_i), p_{RM}(\Delta t)) \tag{S16}$$

Then, the probability of the number $m$ of mutations (between 1 and $L = 22$) is given by:

$$p_m(\mu\Delta t, L) = \frac{\binom{L}{m} (\mu\Delta t)^m (1-\mu\Delta t)^{L-m}}{\sum_{m'=1}^{L} \binom{L}{m'} (\mu\Delta t)^{m'} (1-\mu\Delta t)^{L-m'}} \ , \tag{S17}$$

We sample $m$ positions uniformly from the set of 22 target positions. For each chosen position $i$, the VR nucleotide at that position, $VR_i$, gets replaced by a new nucleotide $q$:

$$q \sim \text{Uniform}\left(\{A, C, G, T\} \setminus \{VR_i\}\right) \tag{S18}$$

The population counts get updated and the second step is executed.

2. Fitness-based growth: see point 2 of scenario S3 A 2

3. Multinomial sampling: see point 3 of scenario S3 A 2

In all 3 scenarios, the population dynamic are simulated for 30 timepoints, using a uniform time-step of $\Delta t = 1$ day. The starting population is common to all scenarios. It is obtained by amplifying the experimental population of Mouse 1 Cage 3 (with ASF, same sample used in S3 A 1), observed at day 3, through a multinomial sampling procedure anologous to that used in the simulations. Specifically, the counts of each $\boldsymbol{VR}$ in the population at $t_1$ are sampled as:

$$(n_{\boldsymbol{VR}^{(1)}}(t_1), \ldots, n_{\boldsymbol{VR}^{(n)}}(t_1)) \sim \text{Multinomial}(N; f_{\boldsymbol{VR}^{(1)}}(t_1), \ldots, f_{\boldsymbol{VR}^{(n)}}(t_1)) \tag{S19}$$

where $f_{\boldsymbol{VR}}(t_1) = \frac{n_{\boldsymbol{VR}}(t_1)}{n(t_1)}$ denotes the frequency of the sequence $\boldsymbol{VR}$ at day $t_1 = 3$, $n(t_1)$ is the total population size (sampled) at day 3 in the experiment, and $N = 10^8$ the total population size in the population dynamics simulation. This value of $N$ is used for the final multinomial sampling, described in point 3, at all time steps for all the scenarios, and has been chosen according to the size of the Bacteroides community gavaged to the mice in the experimental procedure (see Ref. [S1] for the detailed procedure).

###### 3. *Estimation of the average fitness of mutants*

For all scenarios, at each time point $t_i$, the population of mutants is extracted from the total population and a sampling procedure is applied to retrieve a total population size comparable with the experimental one. For the time points at which experimental data are available, $n(t_i)$ is set to the observed total population size, namely:

- $t_i = 3 \to n(3) = 50460$
- $t_i = 7 \to n(7) = 66011$
- $t_i = 10 \to n(10) = 67543$
- $t_i = 14 \to n(14) = 76713$

For the remaining time points of the simulation, $n(t_i)$ is chosen to be equal to the largest size of mutant population experimentally observed (in this case, $n(t_i) = 76713$ for $t_i$ different from 3, 7, 10 and 14 days). The average fitness of the mutated population (VRs different from the parental one noted $\boldsymbol{VR}^{(0)}$) is then calculated as:

$$\langle S_{mut}(t_i)\rangle = \frac{\sum_{\boldsymbol{VR}\neq\boldsymbol{VR}^{(0)}} S(\boldsymbol{VR})\; n_{\boldsymbol{VR}}(t_i)}{\sum_{\boldsymbol{VR}\neq\boldsymbol{VR}^{(0)}} n_{\boldsymbol{VR}}(t_i)} \tag{S20}$$

where $S(\boldsymbol{VR})$ is computed according to (S6).

The whole process is repeated 100 times; for each time point $t_i$, the mean value of $\langle S_{mut}(t_i)\rangle$ across runs, together with its standard deviation, is then plotted.

#### S4. FAST DYNAMICS OF THE VR POPULATION AT FIXED TR

### A. $L = 1$

As detailed in the main text, the population vector $\boldsymbol{n} = (n_q)_{0\le q\le Q-1}$ of the $Q$ nucleotides obeys the matrix equation

$$\frac{\mathrm{d}\boldsymbol{n}}{\mathrm{d}t} = A_{q^*(t)}\boldsymbol{n}\ . \tag{S21}$$

The $A_{q^*(t)}$ matrix corresponds to $\left(A_{q^*(t)}\right)_{q_1,q_2} = \delta_{q_1,q_2}(s\delta_{q_1,q^*(t)} - \nu) + \nu/Q$: all states mutate equally into any other at rate $\nu$, and only the nucleotide $q^*(t)$ (which changes every time $\tau$) has a growth rate of $s$.

The solution is formally expressed as $\boldsymbol{n}(Q\tau) = \prod_{q=0}^{Q-1} \exp\left(\tau A_q\right)\boldsymbol{n}(0) = M\boldsymbol{n}(0)$. At large times $T = kQ\tau$, the population has been through $k$ growth cycles, and the population has been enriched by a factor $M^k$. To get the growth rate of the population under these conditions, we find the largest eigenvalue $\Lambda$ of $M$, which gives the effective population growth rate $S_1(\nu)$ such that

$$e^{S_1(\nu)T} = e^{S_1(\nu)k\tau Q} \approx \sum_p (M^k \cdot \boldsymbol{n}(0))_p \propto \Lambda^k \tag{S22}$$

leading to

$$S_1 = \frac{\ln \Lambda}{Q\tau} \tag{S23}$$

as explained in the main text.

## B. $L > 1$

##### 1. $Q = 2$: perturbation theory

In this section we present the method for deriving the large $L$ effective fitness of a DGR system of length $L$, replacement rate $\nu$ and $Q = 2$ nucleotides (let's note them 0 and 1).
During the time interval $[0, \tau]$, the system selects for 0s: the fitness/growth rate of a sequence with $k$ zeros is simply $sk$. Similarly, during the time interval $[\tau, 2\tau]$, the system selects for 1s: the fitness/growth rate of a sequence with $k$ 0s (and $L-k$ 1s) is simply $s(L-k)$. Then due to the DGR system, each sequence is replaced at rate $\nu$ by a new one, uniformly distributed among any of the $2^L$ possible ones. We fix the scale of time with $s = 1$. The system is cyclic of period $2\tau$.
We are interested in the effective growth rate of the system: if we note $A_0$ (resp. $A_1$) the transition matrix during the first time interval (resp. the second one), then the evolution of the population vector $\boldsymbol{n}$ of size $2^L$ indexed by the sequence (e.g. 11000 for $L = 5$) is given by

$$\boldsymbol{n}(2\tau) = \exp(\tau A_1) \cdot \exp(\tau A_0) \cdot \boldsymbol{n}(0) \, . \tag{S24}$$

Thus, if we note $\Lambda_L$ the dominant eigenvalue of $\exp(\tau A_1) \cdot \exp(\tau A_0)$, the effective growth rate $S_L(\nu)$ of the population is given by:

$$S_L(\nu) = \frac{\ln \Lambda_L}{2\tau} \, . \tag{S25}$$

The objective is to compute $S_L(\nu)$.

*Reduction of the matrix size/Removal of degeneracy.* At first, let's consider the exact form of $A_0$ and $A_1$. They are both $2^L$ square matrices,

$$A_0 = \mathrm{diag}\,(s(\#\text{ of }0) \ - \nu) + \frac{\nu}{2^L}\mathbf{1}\cdot\mathbf{1}^\dagger = D_0 + \frac{\nu}{2^L}\mathbf{1}\cdot\mathbf{1}^\dagger \tag{S26}$$
$$A_1 = \mathrm{diag}\,(s(\#\text{ of }1) - \nu) + \frac{\nu}{2^L}\mathbf{1}\cdot\mathbf{1}^\dagger = D_1 + \frac{\nu}{2^L}\mathbf{1}\cdot\mathbf{1}^\dagger \tag{S27}$$

where $\mathbf{1}$ is the vector with only 1s as entries. A trivial basis of this space is given by the $2^L$ vectors indexed by their ordered sequence of 0s and 1s, that we can note $e_{\boldsymbol{\sigma}}$, $\boldsymbol{\sigma}$ being a vector of size $L$ of 0s and 1s.
We search for eigenvalues of the $A_i$'s. Let us note $\mathcal{E}_k$ the subspace generated by vectors where $k$ nucleotides are 0 (generated by the $e_{\boldsymbol{\sigma}}$'s where $\sum_i \sigma_i = L-k$), and $\mathcal{H}_k = \mathcal{E}_k \cap \{v | v \cdot \mathbf{1} = 0\}$. We note that the dimension of $\mathcal{E}_k$ is $\binom{L}{k}$ while the one of $\mathcal{H}_k$ is $\binom{L}{k} - 1$. Besides, all elements in $v$ in $\mathcal{H}_k$ are eigenvectors of both $A_0$ and $A_1$ of eigenvalues $k$ and $L-k$ respectively:

$$A_0 \cdot v = D_0 \cdot v + \frac{\nu}{2^L}\mathbf{1}\cdot\mathbf{1}^\dagger \cdot v = kv \tag{S28}$$
$$A_1 \cdot v = (L-k)v \tag{S29}$$

Consequently, we found $2^L - (L+1)$ (independent) eigenvectors of $\exp(\tau A_1) \cdot \exp(\tau A_0)$ of same eigenvalue $e^{L\tau}$ as for any $k$ and $v \in \mathcal{H}_k$

$$\exp(\tau A_1) \cdot \exp(\tau A_0) \cdot v = e^{\tau(L-k)+\tau k} v = e^{L\tau} v \, . \tag{S30}$$

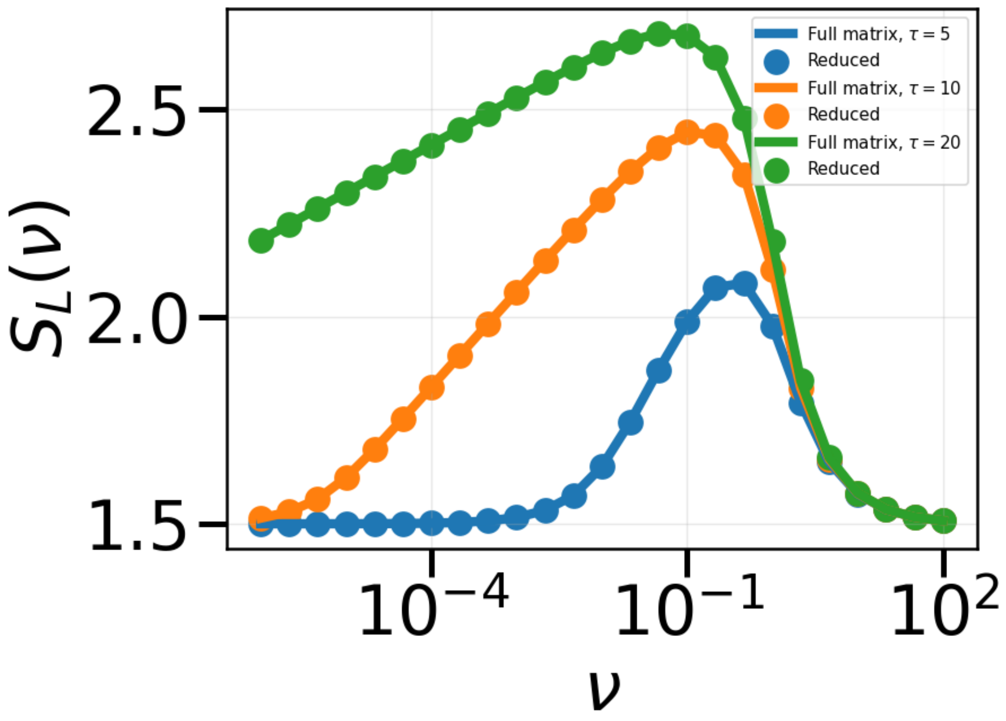


Fig. S2. Comparison of the effective growth rate $S_L(\nu)$ obtained from the top eigenvalue of the $2^L$ matrix and the $L+1$ matrix for $L=3$.

Thus, $\Lambda_L$ is given by the maximum between $e^{L\tau}$ and the maximum eigenvalue on the projected space of dimension $L+1$ corresponding to the remaining eigenvectors to find. An orthonormal basis of this space is simply given by the $L+1$ vectors we will note $|k\rangle$. Each $|k\rangle$ corresponds to the sum of all $\binom{L}{k}$ basis vector in $\mathcal{E}_k$ (vectors with a 1 at the corresponding sequence with $k$ 0s, 0 for all other entries), normalised by $1/\sqrt{\binom{L}{k}}$,

$$|k\rangle = \frac{1}{\sqrt{\binom{L}{k}}} \sum_{\boldsymbol{\sigma} | \sum_i \sigma_i = L-k} e_{\boldsymbol{\sigma}} \ . \tag{S31}$$

By projecting $A_0$ and $A_1$ in this basis, we get the projected $A_0$ and $A_1$ (we keep the same notation in the projected space):

$$A_0 = \sum_{k=0}^{L} (k-\nu)|k\rangle\langle k| + \frac{\nu}{2^L} \sum_{k,\ell=0}^{L} \sqrt{\binom{L}{k}} \sqrt{\binom{L}{\ell}} |k\rangle\langle \ell| \tag{S32}$$

$$A_1 = \sum_{k=0}^{L} (L-k-\nu)|k\rangle\langle k| + \frac{\nu}{2^L} \sum_{k,\ell=0}^{L} \sqrt{\binom{L}{k}} \sqrt{\binom{L}{\ell}} |k\rangle\langle \ell| \tag{S33}$$

where we used that

$$\mathbf{1} \cdot \mathbf{1}^\dagger |k\rangle = \sum_{\ell} \sqrt{\binom{L}{k}} \sqrt{\binom{L}{\ell}} \langle \ell| \tag{S34}$$

This first step gives us a much smaller matrix to diagonalize ($2^L$ versus $L+1$), and we can check our calculations numerically by computing the top eigenvalue for both matrices.

*Perturbation theory at small $\nu/2^L$* Next, we would like to find an approximate expression of the eigenvalues and eigenvectors of both $A_0$ and $A_1$ in order to compute $\Lambda_L$.
For that, we use perturbation theory at first order in $\nu/2^L$, using that at $\nu = 0$ both $A_0$ and $A_1$ are both diagonal in the basis of $(|k\rangle)_{0\leq k\leq L}$, of non-degenerate eigenvalues. We note $\left(|k^{(0)}\rangle\right)_{0\leq k\leq L}$ and $\left(\lambda_k^{(0)}\right)_{0\leq k\leq L}$ the first order shifted eigenvectors and eigenvalues of $A_0$ (resp. $\left(|k^{(1)}\rangle\right)_{0\leq k\leq L}$ and $\left(\lambda_k^{(1)}\right)_{0\leq k\leq L}$ the first order shifted eigenvectors and eigenvalues of $A_1$). We get that:

$$|k^{(0)}\rangle = |k\rangle + \frac{\nu}{2^L}\sum_{\ell\neq k}\frac{\sqrt{\binom{L}{k}}\sqrt{\binom{L}{\ell}}}{k-\ell}|\ell\rangle \tag{S35}$$

$$\lambda_k^{(0)} = k - \nu + \frac{\nu}{2^L}\binom{L}{k} \tag{S36}$$

and

$$|k^{(1)}\rangle = |k\rangle + \frac{\nu}{2^L}\sum_{\ell\neq k}\frac{\sqrt{\binom{L}{k}}\sqrt{\binom{L}{\ell}}}{\ell-k}|\ell\rangle \tag{S37}$$

$$\lambda_k^{(1)} = L - k - \nu + \frac{\nu}{2^L}\binom{L}{k}\,. \tag{S38}$$

Of note , the two basis of eigenvectors are two (distinct) orthonormal basis at first order in $\nu$. From this, we get the formal expression for $\exp(\tau A_1)\cdot\exp(\tau A_0)$:

$$\exp(\tau A_1)\cdot\exp(\tau A_0) = \sum_{k,\ell} e^{(\lambda_k^{(0)}+\lambda_\ell^{(1)})\tau}|k^{(0)}\rangle\langle k^{(0)}|\ell^{(1)}\rangle\langle\ell^{(1)}| \tag{S39}$$

We can evaluate this expression on all the eigenvectors $|k^{(0)}\rangle$, taking the maximal one as a proxy for $\Lambda_L$ (rigorously: a lower bound).

$$\langle k^{(0)}|\exp(\tau A_1)\cdot\exp(\tau A_0)|k^{(0)}\rangle = e^{\lambda_k^{(0)}\tau}\sum_\ell e^{\lambda_\ell^{(1)}\tau}\langle k^{(0)}|\ell^{(1)}\rangle^2 \tag{S40}$$

The exponential terms are maximized for $k = L$ and $\ell = 0$, which provides us with a (good) lower bound for $\Lambda_L$ (by neglecting all other exponential terms):

$$\Lambda_L \geq e^{\lambda_L^{(0)}\tau}e^{\lambda_0^{(1)}\tau}\langle L^{(0)}|0^{(1)}\rangle^2 = e^{\tau(2L-2\nu+2\nu/2^L)}\left(\frac{2\nu}{2^L L}\right)^2 \tag{S41}$$

Finally, we obtain the approximate lower-bound estimate for $S_L(\nu)$:

$$S_L(\nu) \geq \frac{\ln\max\left(e^{L\tau}, e^{\tau(2L-2\nu+2\nu/2^L)}\left(\frac{2\nu}{2^L L}\right)^2\right)}{2\tau} = \max\left(\frac{L}{2}, L-\nu(1-2^{-L}) + \frac{\ln\left(\frac{2\nu}{2^L L}\right)}{\tau}\right) \tag{S42}$$

We can find the optimal DGR switching rate $\nu^*$ by maximizing this expression over $\nu$ and get

$$\nu^* = \frac{1}{\tau(1-2^{-L})} \tag{S43}$$

which confirms the optimality of taking $\nu \approx 1/\tau$ even in the DGR setting with length $L > 1$.
We expect this lower bound to work better at:

- $\nu \ll 1$: because it is perturbation theory in $\nu$.
- $\frac{\nu}{2^L L}e^{\tau L} \geq 1$: the time for best sequence to appear and dominate the population. Otherwise, the exponentials are not dominant and do not dictate the term to keep in the sum. The interpretation is the following: the population currently selected needs some minimal time to grow and dominate by the end of the time window. Thus, the lower bound is poor at $\tau$, $\nu$ and $L$ too small.

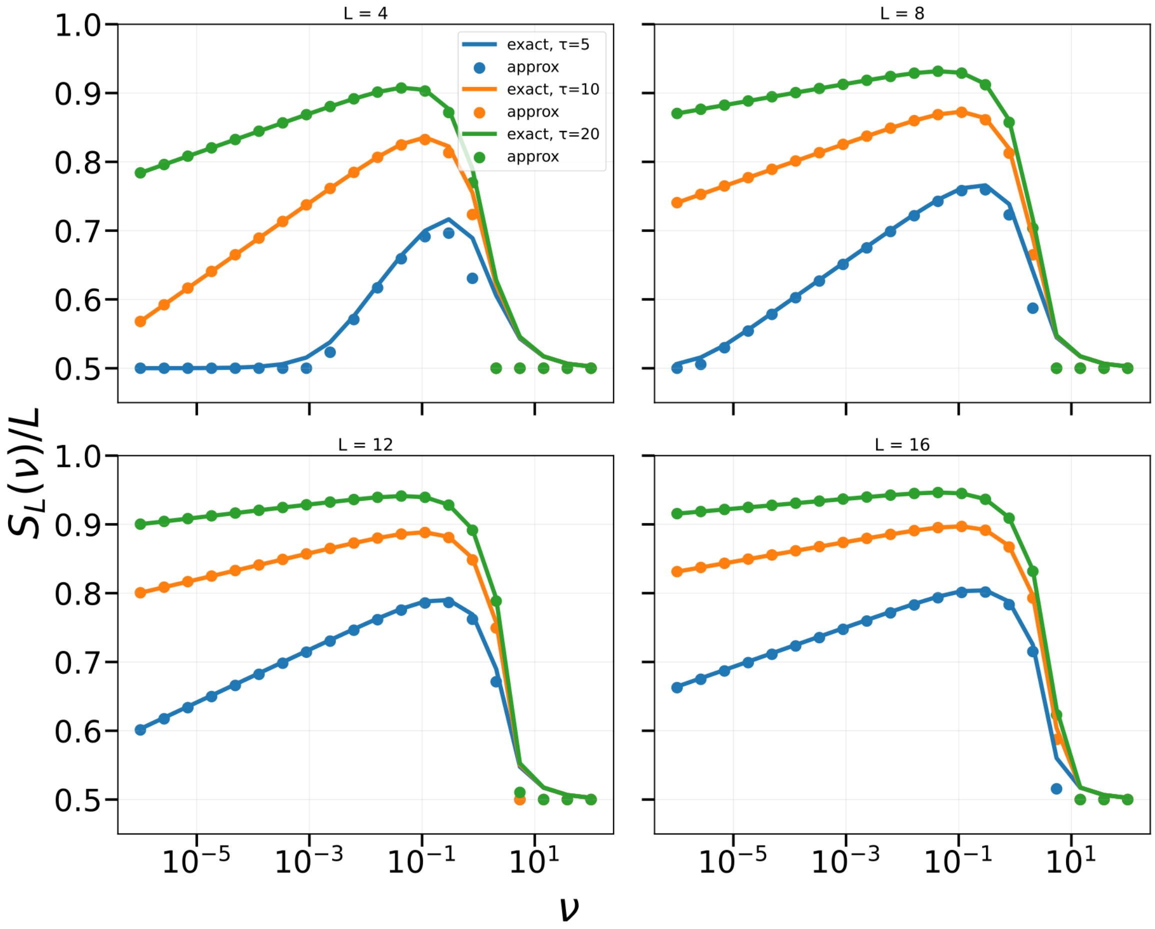


Fig. S3. Lower bound of $S_L(\nu)$ versus its exact value by direct diagonalization of the $L+1$ matrix for $L=4$, $L=8$, $L=12$ and $L=16$.

*Interpretation of the result* At this point, we notice that the result Eq. (S42) can be derived in a simpler way. Let us consider an initial population $\boldsymbol{n}(0)$. During the first time interval $[0,\tau]$, the population with $L$ 0s on their VR noted $n_{0\dots0}$ grows exponentially at a rate $L-\nu+\nu/2^L$, and thus is the dominant population (the total population is approximatively given by $n_{0\dots0}$). In particular, if we also consider the population $n_{1\dots1}$ with $L$ 1s in that time window, their dynamics obey

$$\dot{n}_{1\dots1}(t) = \frac{\nu}{2^L} n_{0\dots0}(t) \tag{S44}$$

such that at time $\tau$,

$$n_{1\dots1}(\tau) \approx \frac{\nu}{L2^L} n_{0\dots0}(\tau) \tag{S45}$$

using that $n_{0\dots0}$ grows exponentially at rate $L-\nu+\nu/2^L \approx L$.
When the environment changes to select 1s, we consider the population of $L$ 0s in their **VR** which switches to $L$ 1s. Besides, we only consider this transition, neglecting all others (individuals which do other transitions are lost). The dynamics is then given by:

$$\dot{n}_{1\dots1}(t) = (L-\nu+\nu/2^L) n_{1\dots1}(t) + \frac{\nu}{2^L} n_{0\dots0}(t)\ . \tag{S46}$$

Because $n_{0\ldots0}(t) = n_{0\ldots0}(\tau)$ for $t > \tau$ (only 1s are selected, 0s have zero growth rate), we have that:

$$n_{1\ldots1}(2\tau) = \frac{\nu}{2^L(L-\nu(1-1/2^L))} n_{0\ldots0}(\tau)\left(e^{(L-\nu+\nu/2^L)\tau}-1\right) + n_{1\ldots1}(\tau)e^{(L-\nu+\nu/2^L)\tau} \tag{S47}$$

$$\approx e^{(L-\nu+\nu/2^L)\tau}\left(\frac{\nu}{2^L L} n_{0\ldots0}(\tau) + n_{1\ldots1}(\tau)\right) \tag{S48}$$

$$\approx e^{(L-\nu+\nu/2^L)\tau}\frac{2\nu}{2^L L} n_{0\ldots0}(\tau)\ . \tag{S49}$$

To summarize, during each environment window, the total population will grow by a factor $\frac{2\nu}{2^L L}e^{(L-\nu+\nu/2^L)\tau}$ at least thanks to the transition $0\ldots0$ to $1\ldots1$. Thus, the effective growth rate is at least

$$S_L(\nu) \geq \frac{\ln\left(\frac{2\nu}{2^L L}e^{(L-\nu+\nu/2^L)\tau}\right)}{\tau} \tag{S50}$$

which gives exactly Eq. (S42) obtained via perturbation theory (the other lower bound $L/2$ coming simply from a population without any DGR switching which grows of a factor $L\tau$ during one bout and stays fixed during the next one). Thus, it means that, to first order in $\nu$, the main contribution of the DGR mechanism to the fitness is given by this transition of the fittest sequence of the previous bout to the next fittest sequence. We will use that directly when considering the case $Q > 2$.

##### 2. General $Q$ lower bound

*Reduction of the matrix size.* We note $A_\kappa$ the rate matrix for state $\kappa$,

$$A_\kappa = \mathrm{diag}\left(s(\#\text{ of }\kappa) - \nu\right) + \frac{\nu}{2^L}\mathbf{1}\cdot\mathbf{1}^\dagger\ . \tag{S51}$$

We can apply the same projection procedure to the $A_\kappa$ matrices to find the highest eigenvalue in the smaller vector space. For any $\kappa$ we have the $A_\kappa$ matrix (similarly to the previous part) for $\kappa = 0, \ldots, Q-1$:

$$A_\kappa = \sum_{c_0,\ldots,c_{Q-1}\mid\sum_k c_k = L} (c_\kappa - \nu)|c_0,\ldots,c_{Q-1}\rangle\langle c_0,\ldots,c_{Q-1}| + \frac{\nu}{Q^L}\sum^{L}_{c_0,\ldots,c_{Q-1},c'_0,\ldots,c'_{Q-1}\mid\sum_k c_k=L,\sum_k c'_k=L}\sqrt{\binom{L}{c_0,\ldots,c_{Q-1}}}\sqrt{\binom{L}{c_0,\ldots,c_{Q-1}}}|c_0,\ldots,c_{Q-1}\rangle\langle c'_0,\ldots,c'_{Q-1}| \tag{S52}$$

where $c_k$ is the number of symbol $k$ in the sequence. We see indeed that this is exact in Fig. S4, meaning that we can use this smaller matrix to compare our approximated results that we will obtain in the following section.

*Effective fitness expression.* A lower bound is given by

$$S_L(\nu) \geq \frac{\ln\max\left(e^{L\tau}, e^{Q\tau(L-\nu+\nu/Q^L)}\left(\frac{2\nu}{Q^L L}\right)^Q\right)}{Q\tau} = \max\left(\frac{L}{Q}, L-\nu(1-Q^{-L}) + \frac{\ln\left(\frac{2\nu}{Q^L L}\right)}{\tau}\right)\ . \tag{S53}$$

It corresponds to the path of states $0\ldots0$ to $1\ldots1$ to $2\ldots2$, so on and so forth where transition from one state to the other happens at rate $\nu/Q^L$. Indeed, because the growth rate is $L$ in the new state, we get that the population $n_1$ in the newly selected state at time $t$ knowing the population in the previously selected state $n_0$ (which will not grow anymore as it is not selected anymore) obeys the equation:

$$\frac{\mathrm{d}n_1}{\mathrm{d}t} = (L-\nu(1-1/Q^L))n_1 + \frac{\nu}{Q^L}n_0 \tag{S54}$$

of solution

$$n_1(\tau) \approx e^{(L-\nu(1-1/Q^L))\tau}\left(\frac{\nu}{LQ^L}n_0(0) + n_1(0)\right) \tag{S55}$$

at large $\tau$. Using that during the previous environment the population $n_0$ grew exponentially at rate $L$ and mutated at rate $\nu/Q^L$ into the population $n_1$, we get as in (S45) that

$$n_1(0) = \frac{\nu}{LQ^L} n_0(0) \ . \tag{S56}$$

which leads to

$$n_1(\tau) \approx e^{(L-\nu(1-1/Q^L))\tau} \frac{2\nu}{Q^L L} n_0(0) \ . \tag{S57}$$

Doing the operation $Q$ times gives us the lower bound above, that we check in Fig. S5. This gives us back $\nu^* = \frac{1}{\tau(1-Q^{-L})}$.

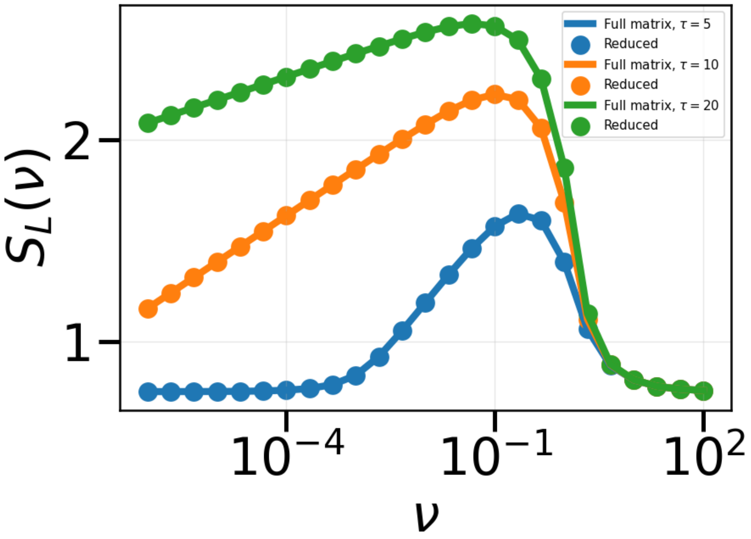


Fig. S4. Comparison of the top eigenvalue of the $Q^L$ matrix and the $O(L^3)$ matrix for $L = 3$.

##### C. Stochastic environment

In this section, we relax the periodicity hypothesis of the enviromental switching. We now draw randomly the new fitness landscape at each time $\tau$: at the switch and at every position, we draw a new nucleotide that will become the new optimal nucleotide for the next time window. In Fig. S6, we show that the asymptotic fitness we obtained previously is in good agreement with the numerical simulations. The difference is more apparent at small $\tau$ and small $L$, as it becomes more frequent to have short time between two selection of the same environment (our calculation neglected non-selected sequences for computation of the effective fitness).

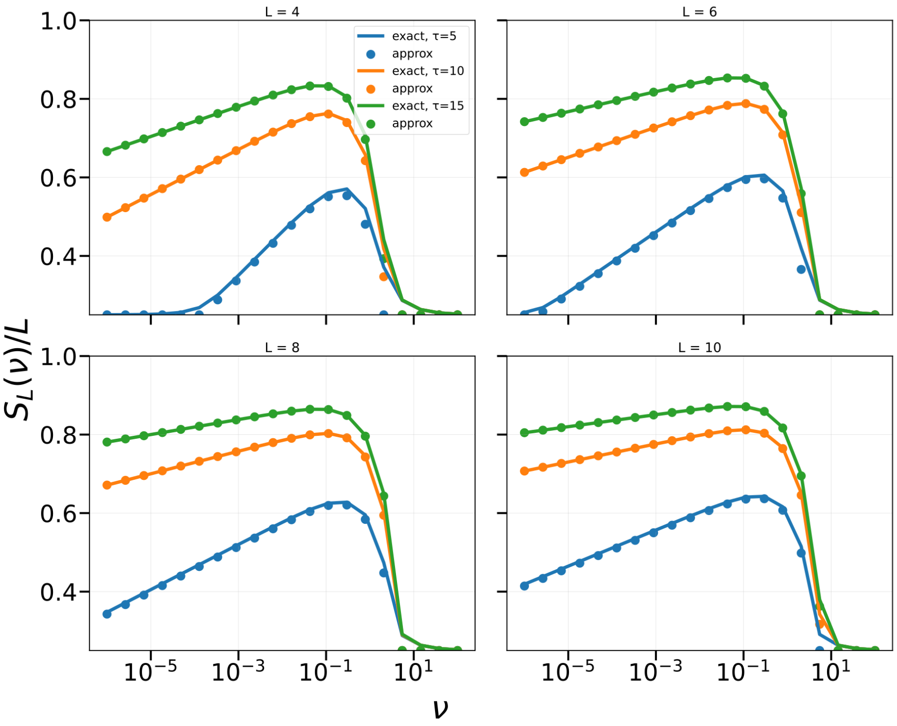


Fig. S5. Lower bound of $S_L(\nu)/L$ versus its exact value by direct diagonalization of the $O(L^3)$ square matrix for $L = 4$, $L = 6$, $L = 8$ and $L = 10$.

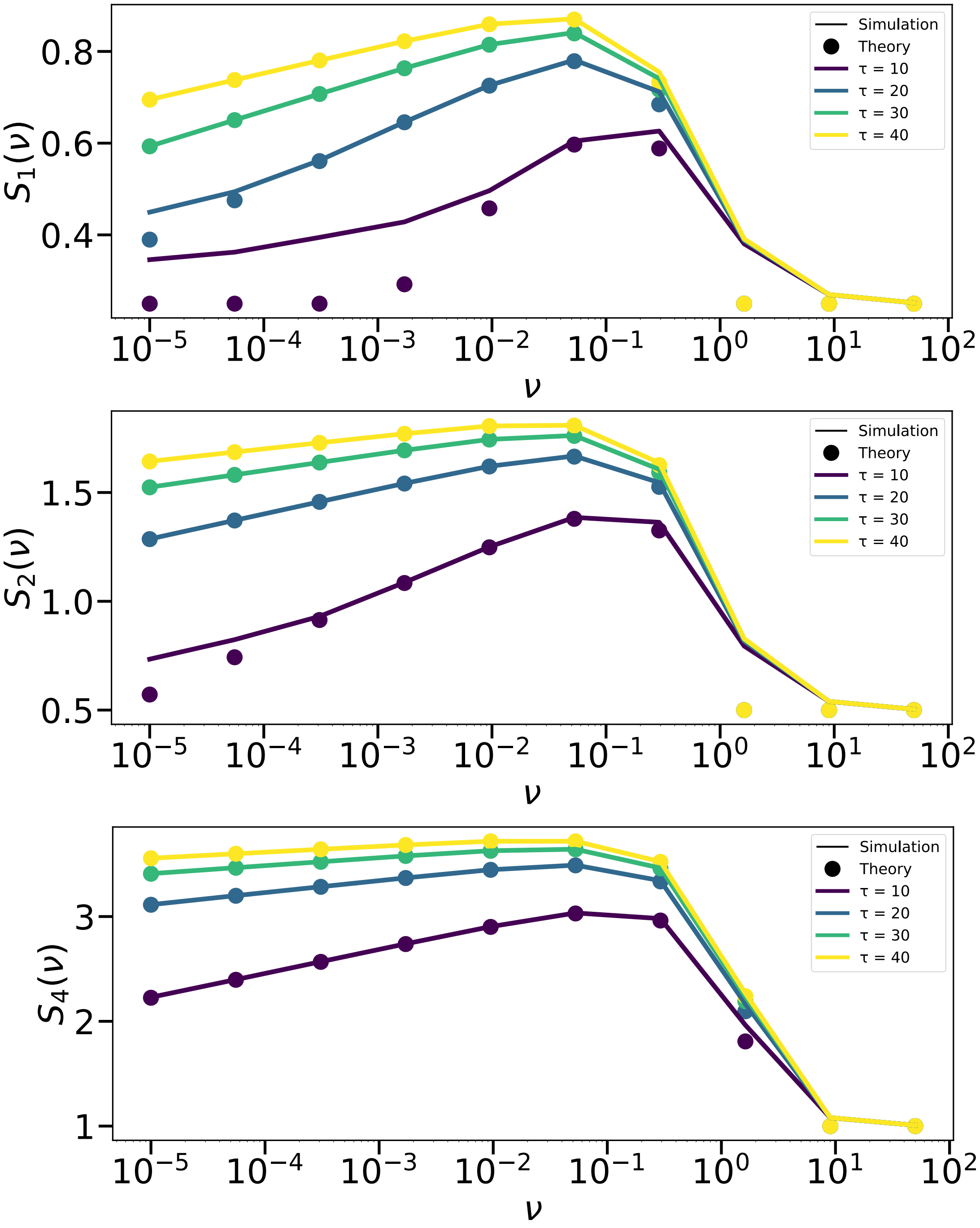


Fig. S6. Asymptotic fitness in the stochastic environment switching. **(a)** L=1 **(b)** L=2 **(c)** L=4.

#### S5. MODEL EXTENSIONS

##### A. On-off DGR

In this subsection, we consider the possibility for the system to activate the DGR diversification at rate $r_+$ and to deactivate it at rate $r_-$. We describe the dynamics by the population vector $(n_{q,\sigma})_{0\leq q\leq Q-1,\sigma=\pm}$ where $+$ denotes the (DGR on) state and $-$ the (DGR) off state. We use the same approach to obtain the asymptotic fitness by describing the transitions between dominant states. We have that $n_{0,-}(t)$ is the dominant population during the previous time window, and $n_{1,-}$ the new population with the fittest genotype. We are interested in the limit of large $\nu$ where the DGR deactivation becomes crucial : because of this, we consider a single DGR on state $n_+$. Indeed, the DGR VR replacement rate is so high that the distribution of VR is uniform along all $Q^L$ possible VRs. Besides, the population has a growth rate of $L/Q$, as for a random VR, there is a probability $1/Q$ that a given nucleotide is the optimal one. The dynamics goes as follow:

$$\frac{\mathrm{d}n_+}{\mathrm{d}t} = \frac{L}{Q}n_+ + r_+n_{1,-} + r_+n_{0,-} - r_-n_+ \tag{S58}$$

$$\frac{\mathrm{d}n_{1,-}}{\mathrm{d}t} = Ln_{1,-} + \frac{r_-}{Q^L}n_+ - r_+n_{1,-} \tag{S59}$$

which results in

$$n_+(t) \approx e^{Lt/Q}\left(n_{0,-}(0)\frac{Qr_+}{L} + n_+(0)\right) - n_{0,-}(0)\frac{Qr_+}{L} \tag{S60}$$

$$n_{1,-}(t) \approx \left[n_{1,-}(0) + \frac{r_-}{Q^LL(1-1/Q)}\left(n_{0,-}(0)\frac{Qr_+}{L} + n_+(0)\right) - \frac{Qr_-r_+}{L^2Q^L}n_{0,-}(0)\right]e^{Lt} \tag{S61}$$

Now, using that in the previous time window, the population $n_{0,-}$ dominated, grew at rate $L$, and transitioned to state $n_+$ at rate $r_+$ while $n_+$ transitioned to $n_{1,-}$ at rate $r_-/Q^L$, we get that:

$$n_+(0) \approx \frac{r_+}{L(1-1/Q)}n_{0,-}(0) \tag{S62}$$

$$n_{1,-}(0) \approx \frac{r_-}{Q^LL}n_+(0) \approx \frac{r_-r_+}{Q^LL^2(1-1/Q)}n_{0,-}(0) \tag{S63}$$

we get that

$$n_{1,-}(\tau) \approx \frac{(2+(1-1/Q)^{-1})r_+r_-}{Q^LL^2(1-1/Q)}e^{L\tau}n_{0,-}(0) \tag{S64}$$

which results in the final growth rate at large $\nu$

$$S_L^{on-off} = \frac{\ln\left(n_{1,-}(\tau)/n_{0,-}(0)\right)}{\tau} \approx L + \frac{\ln\left(\frac{(Q+1+(1-1/Q)^{-1})r_+r_-}{Q^LL^2(1-1/Q)}\right)}{\tau}\,. \tag{S65}$$

We compare the results with numerical simulations in Fig. S7. We observe that despite the addition of the off-on states, the change in the asymptotic fitness is apparent only at large $\nu$, when the DGR system is the most deleterious, and when the population grows faster by using this on-off states (which are not beneficial if $\tau$ is too small).

##### B. Competing populations

In this section, we explicitly model the environment by another population. This framework has been extensively studied using the generalised Lotka-Volterra framework [S5]. We will describe two situations: one with only two genotypes for each population for which we will derive analytical results, and one with extended genotypes for each population, and for which we describe interaction between genotypes via interaction matrices.

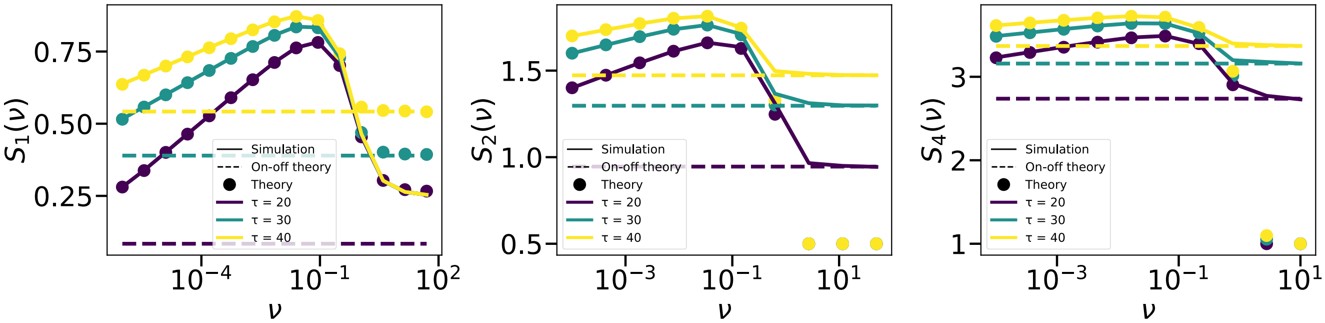


Fig. S7. Asymptotic fitness for DGR with on-off states. $L = 1$ and $L = 2$. $r_- = 10^{-3}$, $r_+ = 10^{-5}$, $N = 10^{12}$, $dt = 0.01$ and the growth rate per base $s = 1$ fixes the timescales. The hashed lines correspond to Eq. S65 while the points correspond to the previous result Eq. S53.

##### *1. Two populations, two genotypes*

We present this for the sake of completeness even though most of the results are more or less known in the litterature [S5]. We note $x(t)$ the frequency of the first population with genotype 1 ($1 - x(t)$ the frequency with genotype 0) and $y(t)$ the frequency of the second population with genotype 1 ($1 - y(t)$ the frequency with genotype 0). We model interactions between the two species by a growth rate $x(t)y(t)$ for the first population with genotype 1 (the first population feeds on the second one if they have the same genotype) while for the second population the growth term is $y(t)(1 - x(t))$ for the population with genotype 1 (the second population escapes the first one).
We add a rate of genotype switching $\nu_x$ and $\nu_y$ for each population. As we will see in the following, we need to consider two time delays $T_x$ and $T_y$ which correspond to the time needed by population $x$ (resp. $y$) to feel changes in the other population. In the end, the dynamics of the populations obey the non-linear equations

$$\begin{cases} \dot{x}(t) = & x(t)y(t-T_x) + \frac{\nu_x}{2}(1-2x(t)) - x(t)\left(x(t)y(t-T_x) + (1-x(t))(1-y(t-T_x))\right) \\ \dot{y}(t) = & sy(t)(1-x(t-T_y)) + \frac{\nu_y}{2}(1-2y(t)) - sy(t)\left((1-y(t))x(t-T_y) + (1-x(t-T_y))y(t)\right) \end{cases} \tag{S66}$$

To understand if one can have oscillations in this system, we study the linear stability of the fixed point $(x^*, y^*) = (1/2, 1/2)$ (the only one when $\nu_x$ and $\nu_y$ are non-zero). By supposing that $(x(t) - x^*, y(t) - y^*) = e^{\lambda t}\boldsymbol{v}$ near the fixed point, we have the matrix equation:

$$\lambda \boldsymbol{v} = \begin{pmatrix} -\nu_x & \frac{e^{-\lambda T_x}}{2} \\ -\frac{se^{-\lambda T_y}}{2} & -\nu_y \end{pmatrix} \boldsymbol{v} \tag{S67}$$

we find $\lambda$ by solving the resolvant equation

$$(\lambda + \nu_x)(\lambda + \nu_y) + \frac{s}{4}e^{-\lambda(T_x+T_y)} = 0 \tag{S68}$$

For $\nu_x = \nu_y = \nu$ and $T = T_x + T_y$, we have an exact expression for $\lambda$:

$$\lambda = \frac{2\,\mathrm{W}_0\left(-\frac{i}{4}e^{\nu T/2}\sqrt{s}\,T\right)}{T} - \nu/2 \tag{S69}$$

where $\mathrm{W}_0$ is the Lambert function. In particular, we can find the critical value of $\nu = \nu_c(T)$ above which the system is not oscillating anymore by solving $\mathrm{Re}\,(\lambda) = 0$. By defining $\lambda = i\omega$, we obtain that

$$\begin{cases} 2\nu\omega - \frac{s}{4}\sin(\omega T) = 0 \\ -\omega^2 + \nu^2 + \frac{s}{4}\cos(\omega T) = 0 \end{cases} \tag{S70}$$

For large times $T$, we suppose that the system's periodicity is large, and note that $\omega T = a$ is fixed. The first line implies that $a = \pi$ as $\nu\omega \to 0$. The second line leads to

$$\nu^2 - \frac{s}{4} = 0 \tag{S71}$$

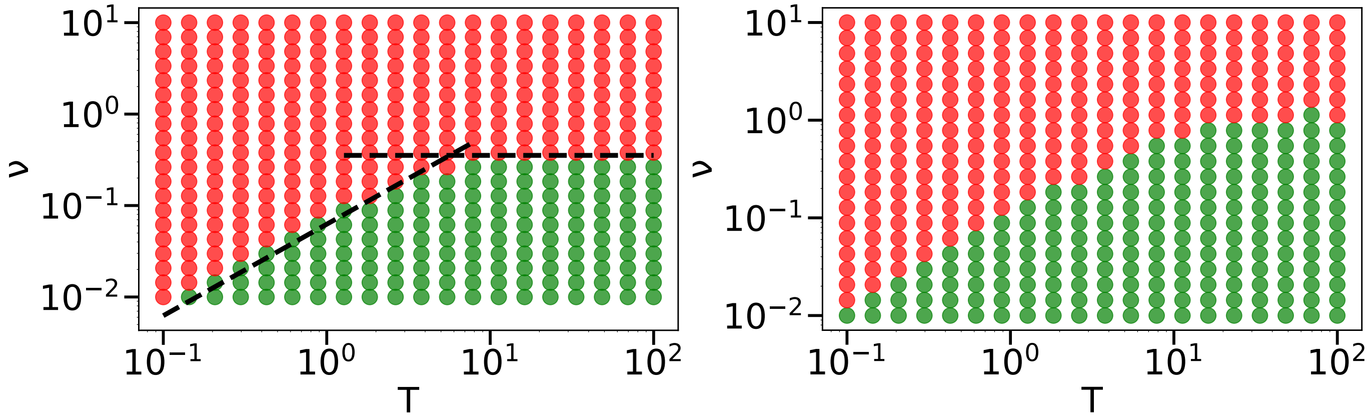


Fig. S8. **Phase diagram of oscillating versus stable competing populations.** In green the points corresponding to oscillating populations in the numerical simulations of Eq. (S66), in red to stable populations. **(a)** $\nu_x = \nu_y = \nu$ and $s = 1/2$. The two regions are separated by the hashed lines corresponding to Eqs. (S72) and (S73). **(b)** $\nu_x = \nu$ and $\nu_y = 0$ and $s = 1/2$.

and finally, for large times $T$, the critical DGR rate is given by

$$\nu_c(T) \sim \frac{\sqrt{s}}{2} \tag{S72}$$

For small times $T$, the periodicity is limited by $\nu$, such that $\omega T \to 0$ and

$$\nu_c(T) \sim \frac{sT}{8} \tag{S73}$$

We represent in Fig. Fig. S8 the regions corresponding to stable and oscillating populations. Additionnally, we show the phase diagram when one population has DGR and the other has not ($\nu_x = \nu > 0$ and $\nu_y = 0$). We see a similar qualitative behaviour, meaning that DGR for one species, regulation of the other by the DGR one and time delayed response is enough to have the periodicity and thus environment fluctuation for the DGR species.

##### 2. *Extended genotypes case*

Here, we consider the situation with explicit modeling of the genotypes interactions. We show that we recover periodicity and oscillating populations even in this more complex situation. If species 1 has length $L$ with $Q$ symbols ($Q = 4$ if nucleotides, $Q = 20$ if amino acids) and species 2 has length $M$ with $Q$ symbols, how to make them interact? A simple idea is to have a matrix $\boldsymbol{W}$ of size $LQ \times MQ$ of interactions. Thus, to evaluate the growth rate of genotype $\vec{\lambda}$ (the one-hot encoded sequence of length $L$ into a 0/1 vector of size $LQ$) versus a genotype $\vec{\mu}$ (the one-hot encoded sequence of length $M$ into a 0/1 vector of size $LQ$), it is given by

$$S_1\left(\vec{\lambda}, \vec{\mu}\right) = \vec{\lambda} \cdot \boldsymbol{W} \cdot \vec{\mu} \tag{S74}$$

while the grotwh rate of genotypes $\vec{\mu}$ is similarly given by the opposite term

$$S_2\left(\vec{\lambda}, \vec{\mu}\right) = -\vec{\lambda} \cdot \boldsymbol{W} \cdot \vec{\mu} \tag{S75}$$

By noting $\boldsymbol{f}$ and $\boldsymbol{g}$ the frequencies of each genotypes (of respectively species 1 and 2), we get the growth rate of genotypes $\vec{\lambda}$ of species 1 in the presence of a population of species 2 described by $\boldsymbol{g}$,

$$S_1\left(\vec{\lambda}\right) = \sum_{\vec{\mu}} \vec{\lambda} \cdot \boldsymbol{W} \cdot \vec{\mu}\, g_{\vec{\mu}} \tag{S76}$$

and similarly for species 2,

$$S_2\left(\vec{\mu}\right) = -\sum_{\vec{\lambda}} \vec{\lambda} \cdot \boldsymbol{W} \cdot \vec{\mu}\, f_{\vec{\lambda}} \tag{S77}$$

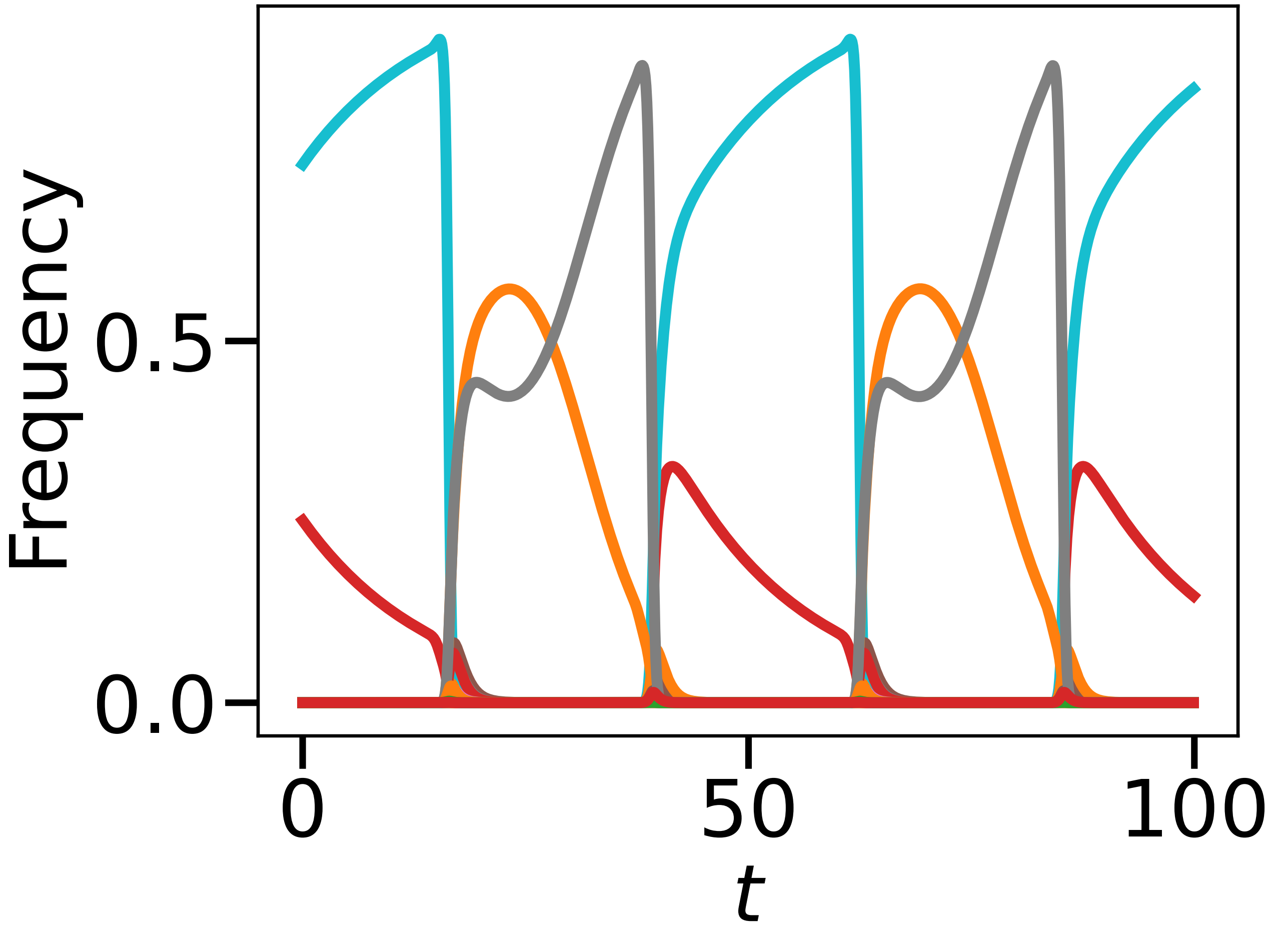


Fig. S9. **Population dynamics under complex genotypes interactions.** We take $Q = 4$ and $L = M = 3$, $W$ coefficients drawn from the uniform distribution in $[-1, 1]$, $\tau = 10$ and $\nu = 10^{-2}$.

Finally, this gives the dynamics of each species

$$\frac{\mathrm{d}f_{\vec{\lambda}}}{\mathrm{d}t} = \left( S_1\left(\vec{\lambda}\right) - \sum_{\vec{\lambda}'} f_{\vec{\lambda}'} S_1\left(\vec{\lambda}'\right) \right) f_{\vec{\lambda}} - \nu f_{\vec{\lambda}} + \frac{\nu}{Q^L} \tag{S78}$$

$$\frac{\mathrm{d}g_{\vec{\mu}}}{\mathrm{d}t} = \left( S_2\left(\vec{\mu}\right) - \sum_{\vec{\mu}'} g_{\vec{\mu}'} S_2\left(\vec{\mu}'\right) \right) g_{\vec{\mu}} - \nu g_{\vec{\mu}} + \frac{\nu}{Q^M} \tag{S79}$$

In Fig. Fig. S9, we represent the dynamics of $x$ genotypes: the population oscillates between a large variety of genotypes without any external driving. This shows that our simple modelisation can be easily extended to account for explicit interactions using the genotypes, while keeping its qualitative properties.

#### C. Epistatic landscape: single genotype selection

##### 1. Single genotype selected

In this section, we consider the most extreme (and simplest) case of epistasis: there is a single genotype with maximum fitness advantage $s$, while all the others present no advantage (growth rate 0). In other words, in this model, the fitness $S(\boldsymbol{VR}, t)$ is given

$$S(\boldsymbol{VR}, t) = s\delta_{\boldsymbol{VR}, \boldsymbol{VR}^*(t)} \tag{S80}$$

According to the calculations made for the DGRs in the additive fitness framework, the main contribution to the fitness comes from the DGR event that connects the previous optimum sequence to the new one. Thus, we expect that the approximation of the effective fitness $S_L(\nu)$ to remain valid in this new framework, highlighting DGR resilience to epistasis (at high $\nu$ however, the downward limit becomes $s/Q^L$ as now, only $1/Q^L$ of the sequences have fitness gain).
However, when considering par base mutation at rate $\mu$, the epistasis effect can be deleterious: the system needs to do $L$ mutations without any fitness gain before reaching the optimum sequence. We expect a strong decrease of fitness in this situation. We explain below how to compute the effective fitness for this system.
By defining $n_k(t)$ the population size of sequences with $k$ nucleotides corresponding to the optimum sequence, we have the following system of equations (with convention $n_{-1} = n_{L+1} = 0$):

$$\dot{n}_k(t) = \left( s\delta_{k,L} - \frac{\mu[(L-k) + (Q-1)k]}{Q} \right) n_k(t) + \frac{\mu(L-k+1)}{Q} n_{k-1}(t) + \frac{\mu(Q-1)(k+1)}{Q} n_{k+1}(t) \tag{S81}$$

Once again, we can find an approximate formula by considering the path contributing the most to the fitness growth, $0 \to 1 \to \ldots \to L$, such that $(n_k(t))_{0 \leq k \leq L}$ obeys a triangular equation we can solve.

$$n_k(t) = \sum_{i=0}^{k} n_i(0) \prod_{m=i+1}^{k} \frac{\mu(L-m+1)}{Q} \sum_{j=i}^{k} e^{\lambda_j t} \prod_{m=i, m\neq j}^{k} \frac{1}{\lambda_j - \lambda_m} \tag{S82}$$

$$\text{with} \quad \lambda_j = s\delta_{j,L} - \frac{\mu}{Q}\big(L + (Q-2)j\big) \tag{S83}$$

To have the $n_i(0)$'s as a function of $n_0(0)$, the dominating population at time $t = 0$, we use that for $t < 0$, $n_0(0)$ grows at rate $s$, and solve the same Eq. (S81) replacing $\delta_{k,L}$ by $\delta_{k,0}$:

$$n_i(0) \approx \left( \frac{\mu}{Qs} \right)^i \frac{L!}{(L-i)!} n_0(0) \tag{S84}$$

which finally gives

$$n_L(t) \approx \left( \frac{\mu}{Qs} \right)^L (L+1)! n_0(0) \tag{S85}$$

Thus, using $S_L(\mu) \approx \ln\left(n_L(\tau)/n_0(0)\right)/\tau$ is enough to compute the approximate effective fitness. In Fig. Fig. S10, we compare the initial DGR formula with the simulation in the epistatic environment, as well as the effective fitness for the per base mutation mechanism at rate $\mu$ , $S_L(\mu)$, all in $s = L$ case. A good approximation of the effective fitness is thus given by

$$S_L(\mu) \approx \frac{1}{\tau} \ln\left( \frac{(L+1)!\mu^L}{Q^L s^L} \right) + s - \frac{\mu(Q-1)L}{Q} \tag{S86}$$

We can compare the result with the one for DGR: the log cost factor comes with a $(\mu/s)^L$ term while in the DGR it was $\nu/s$. It is linearly more expensive not to have DGR in this fully epistatic landscape, a result which goes in the direction saying that DGR is most beneficial in situations where the landscape presents strong epistasis.

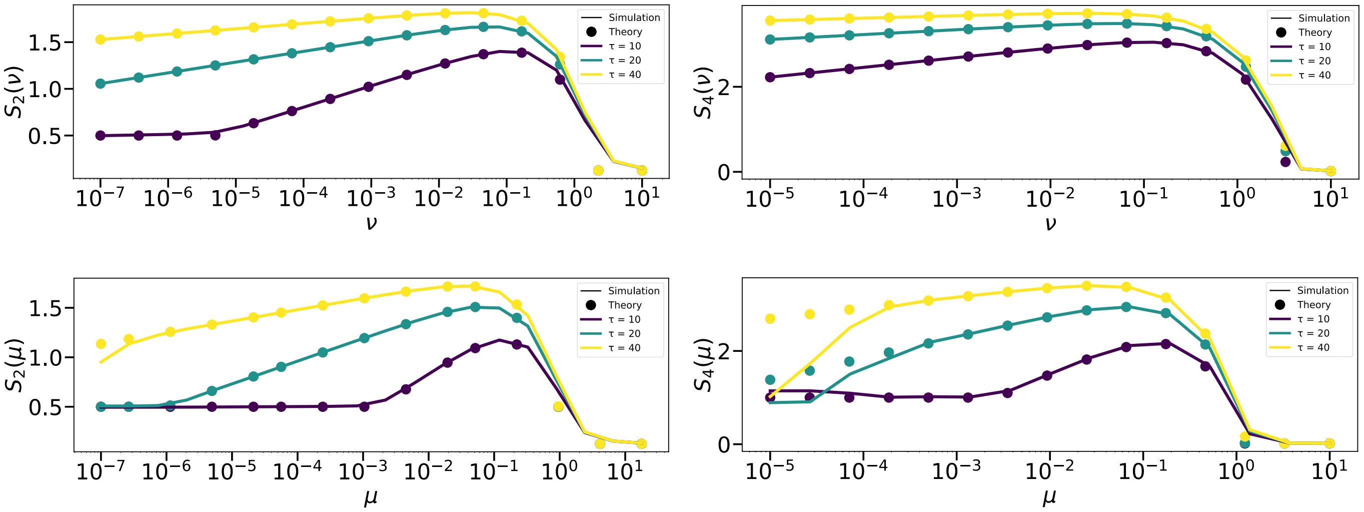


Fig. S10. **Effective Fitness in Epistatic framework.** **(a)** DGR, $L = 2$ **(b)** DGR, $L = 4$ **(c)** Per base mutation, $L = 2$ **(d)** Per base mutation, $L = 4$ .

##### *2. Generalization*

We now consider fitness landscapes which depend only to the distance to the optimal sequence. These landscapes can be considered to be non-linear transformations of the additive landscape. Thus, noting $d(\boldsymbol{VR}, \boldsymbol{VR'})$ the Hamming distance between sequences $\boldsymbol{VR}$ and $\boldsymbol{VR'}$, the fitness $S\left(\boldsymbol{VR}, t\right)$ is given by:

$$S\left(\boldsymbol{VR}, t\right) = \sigma(d(\boldsymbol{VR}, \boldsymbol{VR}^*(t))) \tag{S87}$$

where $\sigma$ is a strictly increasing function. Making the same calculations as in the previous section, we obtain that

$$n_L(t) \approx n_0(0) \left(\frac{\mu}{Q}\right)^L (L+1)! \prod_{i=1}^{L} \frac{1}{\sigma(L) - \sigma(L-i)} \tag{S88}$$

which leads to

$$S_L(\mu) \approx \frac{1}{\tau} \ln \left( \frac{(L+1)!\mu^L}{Q^L} \prod_{i=1}^{L} \frac{1}{\sigma(L) - \sigma(L-i)} \right) + \sigma(L) - \frac{\mu(Q-1)L}{Q} \ . \tag{S89}$$

#### S6. SLOW DYNAMICS OF THE TR

### A. $L = 1$

To describe the **TR** dynamics, we will separate the discussion by separating over the diverse type of **(TR,VR)** genotypes. We note $A$ the nucleotide which corresponds to the diversification (when present in the **TR**), $T$ corresponding to the currently selected nucleotide (when present in the **VR**), and $N$ for all the rest (we note $M$ when we include $A$ in this population). We represent the graph of population fluxes between the different **(TR, VR)** populations.

Of note, we make the assumption that $T \neq A$, hypothesis we discuss in the end of the section.

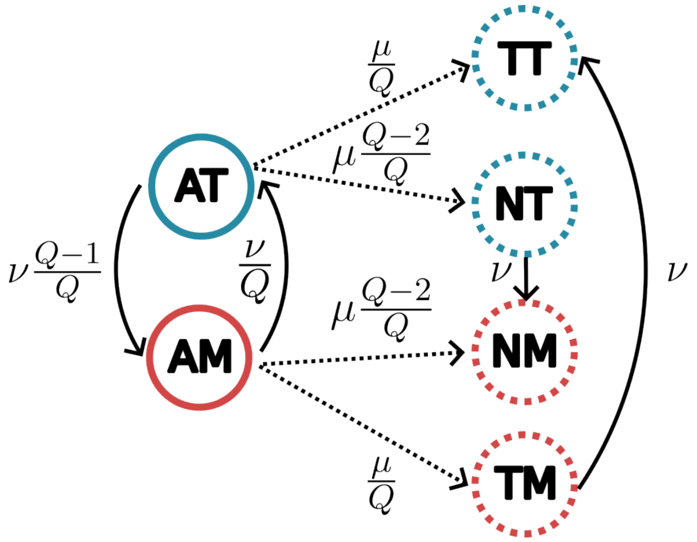


Fig. S11. **The graph of population transitions.** The graph of effective transitions between the different **(TR,VR)** population (first letter is the **TR**, second the **VR**). $A$ corresponds to the diversifying nucleotide, $T$ to the currently selected nucleotide, $M$ to any nucleotide except form $T$, $N$ to any nucleotide except from $A$ and $T$. In blue are the population with growth rate 1, in red those with growth rate 0. Hashed circles correspond to population of size of order $\mu$, while plain lines are of order 1. Dotted arrows represent transitions of order $\mu$, plain line arrows those of order $\nu$. We do not represent transitions which would lead to $\mu^2$ corrections to the population sizes.

###### 1. *The* **TR**=*A population: first order*

As seen in the first part, by taking $\nu \approx 1/\tau$ we are going to select for **TR** with the Adenine. Thus, as a first step, we describe this population, which should dominate the whole population. To do so, we split this population in two: either they have the right **VR**=T (population of size $n_{AT}(t)$) or they don't (population of size $n_{AM}(t)$). The equations describing the joint dynamics of these two populations are given by:

$$\dot{n}_{AT} = \left(1 - \nu\frac{Q-1}{Q}\right) n_{AT} + \frac{\nu}{Q} n_{AM} \tag{S90}$$

$$\dot{n}_{AM} = -\frac{\nu}{Q} n_{AM} + \nu\frac{Q-1}{Q} n_{AT} \tag{S91}$$

of solution given by:

$$n_{AT}(t) = C_+ e^{\lambda_+ t} + C_- e^{\lambda_- t} \tag{S92}$$

$$n_{AM}(t) = C_+ x_+ e^{\lambda_+ t} + C_- x_- e^{\lambda_- t} \tag{S93}$$

where

$$\Delta = \sqrt{(1-\nu)^2 + \frac{4\nu}{Q}}, \qquad \lambda_\pm = \frac{1 - \nu \pm \Delta}{2}. \tag{S94}$$

and

$$x_\pm = \frac{Q\lambda_\pm - Q + \nu(Q-1)}{\nu} \tag{S95}$$

$$C_+ = \frac{n_{AM}(0) - x_+ n_{AT}(0)}{x_+ - x_-} \tag{S96}$$

$$C_- = \frac{x_+ n_{AT}(0) - n_{AM}(0)}{x_+ - x_-} \tag{S97}$$

For the initial conditions, we use that at times $t \gg 1/(\lambda_+ - \lambda_-) = 1/\Delta$,

$$\frac{n_{AM}(t)}{n_{AT}(t) + n_{AM}(t)} \sim \frac{x_+}{1+x_+} \,. \tag{S98}$$

However, we know that the initial fraction of the population in $(A,T)$ and $(A,M)$ are given by the respective fractions at the previous time window, where the $(A,T)$ population corresponds to a fraction $1/(Q-1)$ of the previous $(A,M)$ population ($T$ not being the selected nucleotide in the previous time window and supposing that the behaviour of all non positively selected nucleotide is the same). Thus, the initial fraction of sites starting in state **(TR,VR)**=(A,T) can be approximated by

$$n_{AT}(0) = \frac{x_+}{(Q-1)(1-x_+)} \,, \; n_{AM}(0) = 1 - n_{AT}(0) \tag{S99}$$

From this, we get the fraction of the population in each state,

$$f_{AT}(t) = \frac{n_{AT}(t)}{n_{AM}(t) + n_{AT}(t)} \,, \; f_{AM}(t) = \frac{n_{AM}(t)}{n_{AM}(t) + n_{AT}(t)} \,. \tag{S100}$$

Using this, we get the full description of the **TR**=A population as well as the average fitness $\lambda(t)$ which is given by the fraction of the population in the $AT$ state:

$$\lambda(t) = 1 \times f_{AT}(t) + 0 \times f_{AM}(t) = f_{AT}(t) \tag{S101}$$

We check our analytical results against numerical simulations in Fig. S12.

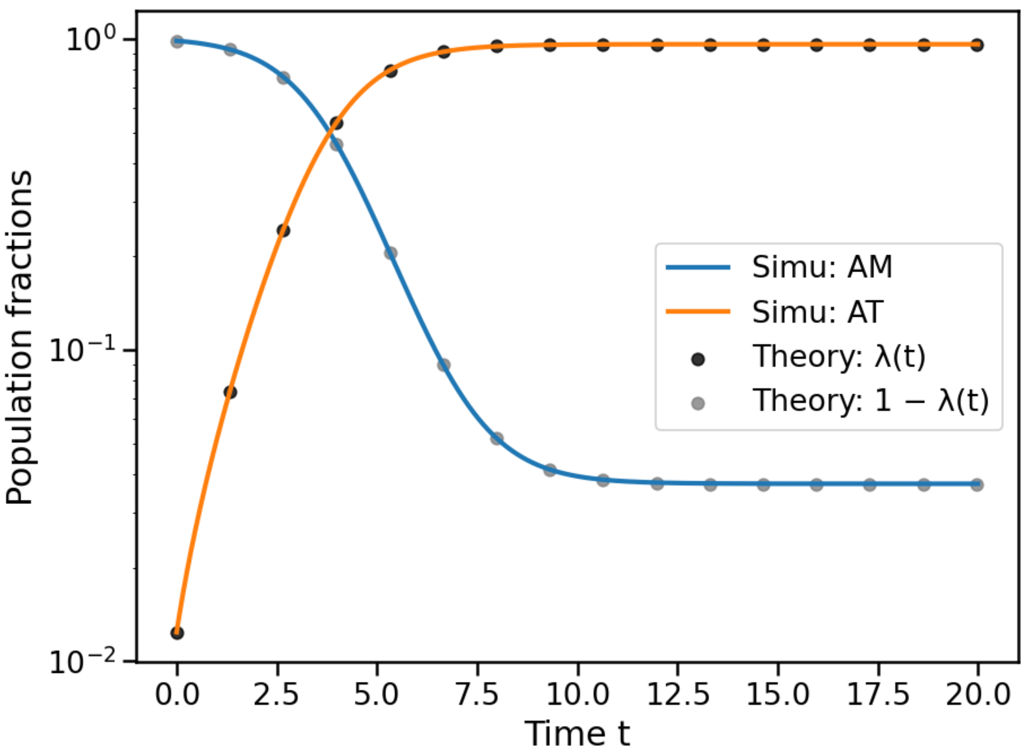


Fig. S12. **A population fraction.** Comparison between numerical simulations (lines) and theoretical predictions (dots) of the population of **TR** having an adenine. The population size is $10^7$, and the parameters are $\tau = 20$, $\nu = 5.10^{-2}$, $\mu = 10^{-5}$, $Q = 4$.

##### *2. The* **TR**$\neq A$ *population: the order* $\mu$ *correction*

To describe this population, we directly compute the fraction dynamics using Wright-Fisher equations [S6] as we computed the average fitness dynamics $\lambda(t)$ (to first order, without the $\mu$ corrections). In the following, we can replace when necessary $f_{AT}(t) = \lambda(t)$ and $f_{AM}(t) = 1 - \lambda(t)$. We will note $\Lambda(t) \equiv \int_0^t \lambda(s)\mathrm{d}s$.

**TR**$=T$, **VR**$= M$

$$\dot{f}_{TM}(t) = (-\nu - \lambda(t))\, f_{TM}(t) + \frac{\mu}{Q} f_{AM}(t) \tag{S102}$$

such that

$$f_{TM}(t) = f_{TM}(0)e^{-\nu t - \Lambda(t)} + \frac{\mu}{Q} e^{-\nu t - \Lambda(t)} \int_0^t e^{\nu s + \Lambda(s)} f_{AM}(s)\mathrm{d}s \tag{S103}$$

$$= f_{TM}(0)e^{-\nu t - \Lambda(t)} + \frac{\mu}{Q} e^{-\nu t - \Lambda(t)} \int_0^t e^{\nu s + \Lambda(s)} (1 - \lambda(s))\mathrm{d}s \tag{S104}$$

**TR**$=T$, **VR**$=T$

$$\dot{f}_{TT}(t) = (1 - \lambda(t)) f_{TT}(t) + \frac{\mu}{Q} f_{AT}(t) + \nu f_{TM}(t) \tag{S105}$$

such that

$$f_{TT}(t) = f_{TT}(0)e^{t - \Lambda(t)} + e^{t - \Lambda(t)} \int_0^t \left( \frac{\mu}{Q} f_{AT}(s) + \nu f_{TM}(s) \right) e^{-s + \Lambda(s)}\mathrm{d}s \tag{S106}$$

$$= f_{TT}(0)e^{t - \Lambda(t)} + e^{t - \Lambda(t)} \int_0^t \left( \frac{\mu}{Q} \lambda(s) + \nu f_{TM}(s) \right) e^{-s + \Lambda(s)}\mathrm{d}s \tag{S107}$$

**TR**=$N$, **VR**=$T$

$$\dot{f}_{NT}(t) = (1-\lambda(t)-\nu)f_{NT}(t) + \frac{\mu(Q-2)}{Q}\left(f_{AT}(t)+f_{TT}(t)\right) \tag{S108}$$

We neglect $f_{TT}(t) \ll f_{AT}(t)$:

$$f_{NT}(t) = f_{NT}(0)e^{t-\Lambda(t)-\nu t} + \frac{\mu(Q-2)}{Q}e^{t-\Lambda(t)-\nu t}\int_0^t e^{-s+\Lambda(s)+\nu s} f_{AT}(s)\mathrm{d}s \tag{S109}$$

$$= f_{NT}(0)e^{t-\Lambda(t)-\nu t} + \frac{\mu(Q-2)}{Q}e^{t-\Lambda(t)-\nu t}\int_0^t e^{-s+\Lambda(s)+\nu s} \lambda(s)\mathrm{d}s \tag{S110}$$

**TR**= $N$, **VR**=$M$

$$\dot{f}_{NM}(t) = -\lambda(t)f_{NM}(t) + \frac{\mu(Q-2)}{Q}f_{AM}(t) + \nu f_{NT}(t) \tag{S111}$$

of solution given by

$$f_{NM}(t) = f_{NM}(0)e^{-\Lambda(t)} + e^{-\Lambda(t)}\int_0^t e^{\Lambda(s)}\left(\frac{\mu(Q-2)}{Q}f_{AM}(s) + \nu f_{NT}(s)\right)\mathrm{d}s \tag{S112}$$

$$= f_{NM}(0)e^{-\Lambda(t)} + e^{-\Lambda(t)}\int_0^t e^{\Lambda(s)}\left(\frac{\mu(Q-2)}{Q}(1-\lambda(s)) + \nu f_{NT}(s)\right)\mathrm{d}s \tag{S113}$$

We then have to solve the self consistent equation which connects the different fractions together:

$$f_{TM}(0) = \frac{f_{NT}(\tau)}{Q-2} \tag{S114}$$

$$f_{TT}(0) = \frac{f_{NM}(\tau)}{Q-2} \tag{S115}$$

$$f_{NT}(0) = f_{TM}(\tau)\frac{Q-2}{Q-1} \tag{S116}$$

$$f_{NM}(0) = f_{TT}(\tau) + f_{TM}(\tau)\frac{Q-2}{Q-1} + f_{NM}(\tau)\frac{1}{Q-2} + f_{NT}(\tau)\frac{1}{Q-2}\ . \tag{S117}$$

The idea behind the self consistent equations is to consider that, at the next time epoch, it will be a different nucleotide (let's say G) that will be selected. We have to consider, when the selected nucleotide changes, where are the different populations going to be splitted.

- TT: going to NM population (not the best nucleotide in new time window).
- NT: a fraction $1/(Q-2)$ of the population is GT (N can be any of the $Q-2$ nucleotides) and thus goes to the GM population. The rest is going to the NM population (neither the selected nucleotide at **TR** or at **VR**).
- TM: a fraction $1/(Q-1)$ of the population is TG and goes to the NG population. The rest goes to the NM population.
- NM: Because of the rapid replacement by the DGR activity, this population is mostly constituted in equal fractions of (**TR**, **VR**) population where the **TR** matches the **VR**. In that case, there are effectively $Q-2$ subpopulations, with one corresponding to the GG population (thus a fraction $1/(Q-2)$). The rest goes to the $NM$ population. The other part of the population where the **TR** does not match the **VR** behaves similarly as the $TM$ population, in particular all the populations where the **VR** is a G and the **TR** is not an $A$, a $T$ or a $G$ (there are $Q-3$ of those): these will be added to the NG population.

At this point, we observe as in the first part of the main text that the **TR**≠A population dynamics can be expressed as a linear problem: indeed, the time evolution between $[0,\tau[$ can be expressed as a linear transformation of the frequencies vector $\vec{f} \to M\vec{f} + \vec{b}$ (the expressions at time $\tau$ being affine in those at time 0), while the final transformation of the subpopulations when the optimum sequence changes is of the form $\vec{f} \to C\vec{f}$. Thus, to find the

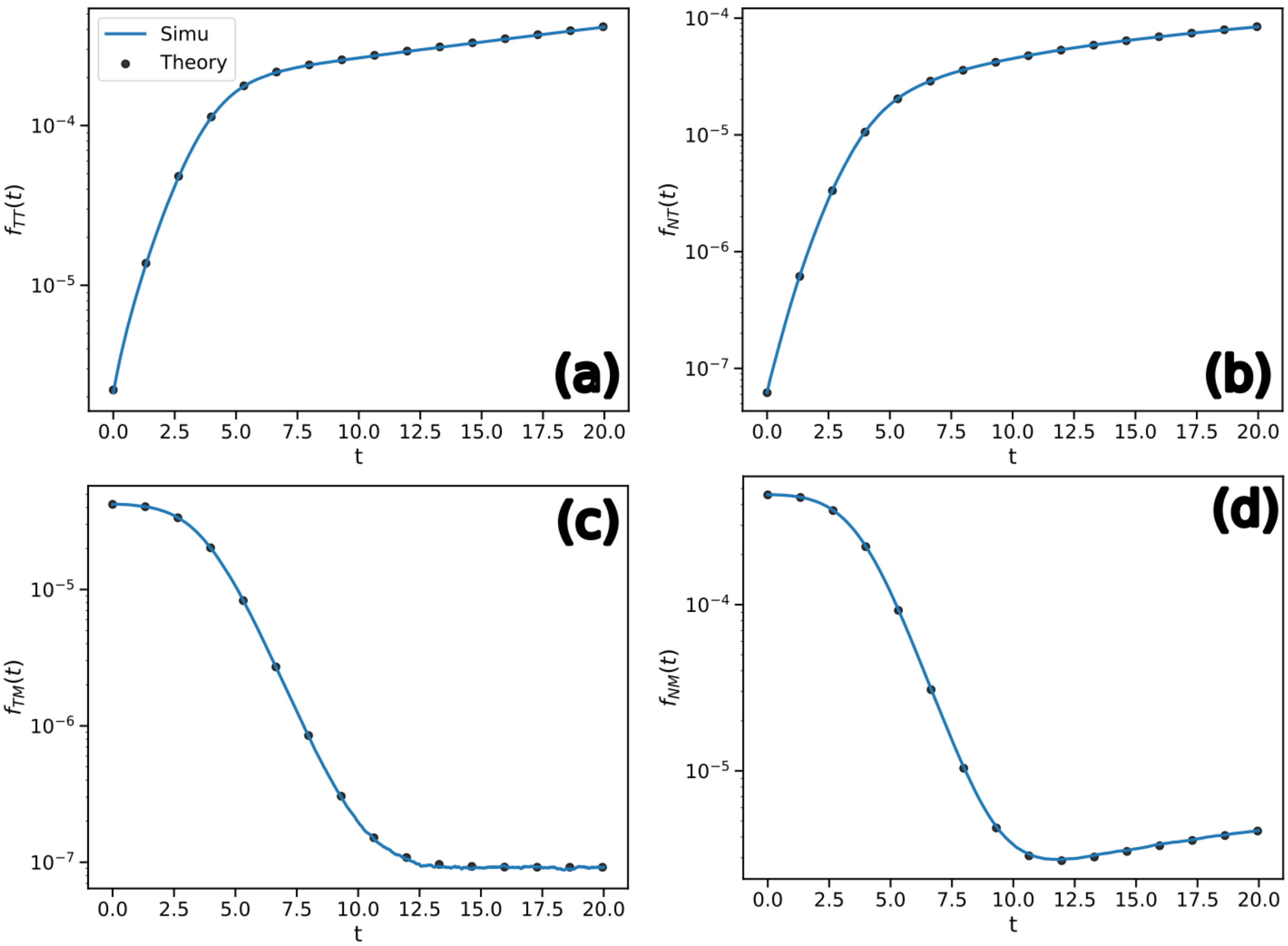


Fig. S13. **Non A populations fraction.** Comparison between numerical simulations (lines) and theoretical predictions (dots) of the fraction of the population of **TR** not having an adenine. **(a)** The **TR**=T, **VR**=T population, **(b)** The **TR**=T, **VR**=M population,**(c)** The **TR**=N, **VR**=T population,**(d)** The **TR**=N, **VR**=M population. The population size is $10^{12}$, and the parameters are $\tau = 20$, $\nu = 5.10^{-2}$, $\mu = 10^{-5}$, $Q = 4$.

initial conditions self-consistently, we have to find the solution $\vec{f}^*$ of the matrix problem $\vec{f} = C \cdot M\vec{f} + C\vec{b}$ .

We check that Eqs. (S114) to (S117) allow to correctly predict the dynamics of each population. In Fig. S13, we use the solution $\vec{f}^* = (I - C \cdot M)^{-1} \cdot C\vec{b}$ at time $\tau$ as initial condition for the results of subsections 1 to 4. We observe a perfect match between our simulations during the time window $[0, \tau]$ and the analytical results. Besides, we confirm that the $TT$ population dominates the population (of **TR**$\neq$A) early in the dynamics, as supposed in the next section text.

From these, we can evaluate the fraction of the population which has for **TR** a nucleotide which is not $A$ as a function of time, displayed in Fig. S14. Finally, we can evaluate the average fraction $f_{\text{not A}}$ sequences as a function of the parameters of the model and compare it to the results of numerical simulations. Once again, we obtain a perfect match with our analytical result.

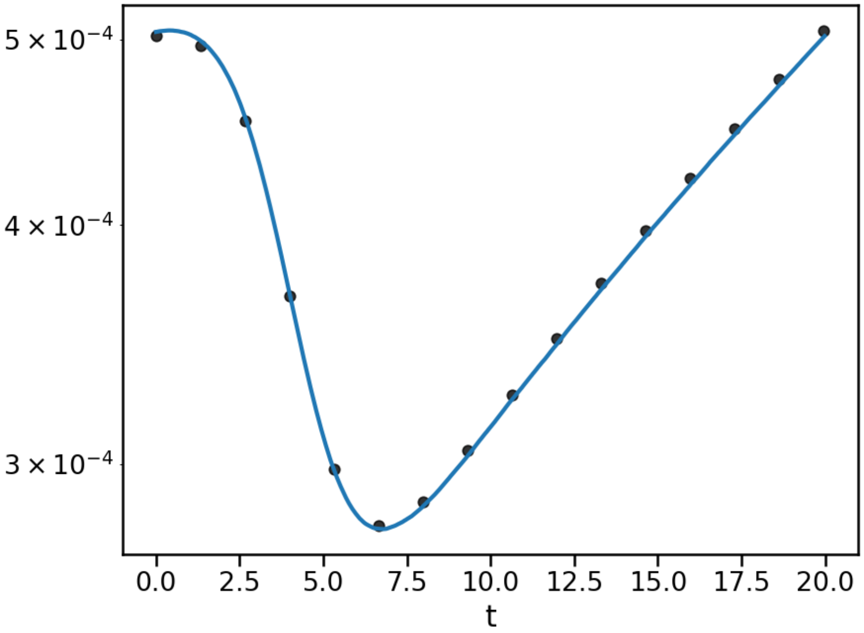


Fig. S14. **Non A population fraction.** Comparison between numerical simulations (lines) and theoretical predictions (dots) of the fraction of the population of **TR** not having an adenine as a function of time. The population size is $10^{12}$, and the parameters are $\tau = 20$, $\nu = 5.10^{-2}$, $\mu = 10^{-5}$, $Q = 4$.

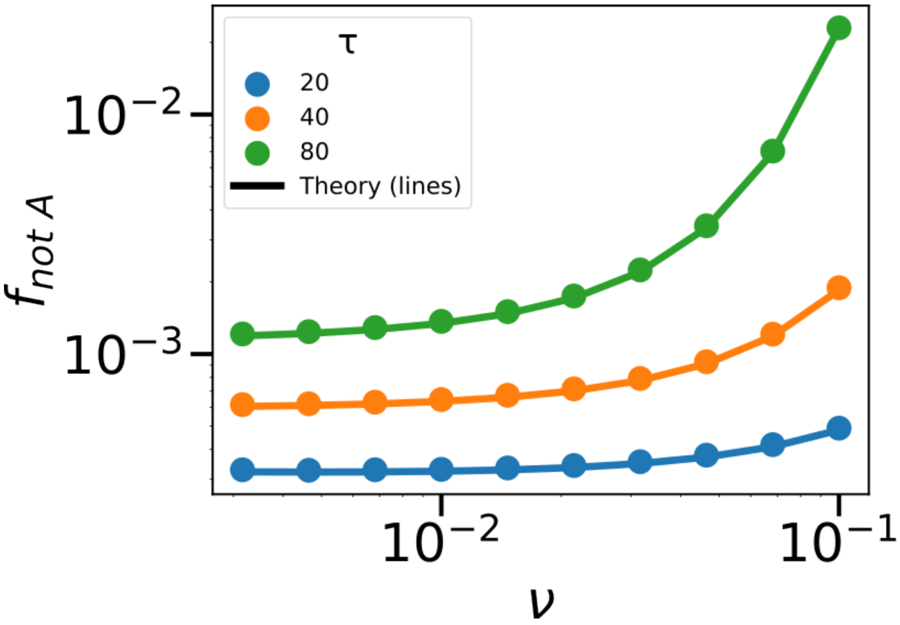


Fig. S15. **Non A population fraction.** Comparison between numerical simulations (lines) and theoretical predictions (dots) of the fraction of the population of **TR** not having an adenine. The population size is $10^{12}$, and the parameters are $\mu = 10^{-5}$, $Q = 4$.

##### 3. Simplified dynamics

We offer to do a simple approximation here to better understand the dynamics of the **TR** population. The idea is that we consider the population with an Adenine in the **TR** as the dominant one. As in the previous sections, we note $f_{AT}(t)$ (resp. $f_{AM}(t)$) the fraction of the population with an $A$ at the **TR** and the currently selected nucleotide (resp. any non-selected nucleotide) noted $T$ (resp. $M$) in the **VR**, with $T \neq A$ otherwise $A$ is surely selected in both the **TR** and the **VR**. The fitness advantage of having $T$ at the **VR** is $s$. In the following, we fix time scales by taking $s = 1$ (which is equivalent as taking the inverse growth rate as the unit of time). We use Wright-Fisher equations [S6] to describe the population dynamics and note $\lambda(t) = 1 \times f_{AT}(t) + 0 \times f_{AM}(t)$ the average population fitness. At first, because $\nu \gg \mu$, we neglect the **TR** mutation rate $\mu$ for the determination of the fraction of the population in each state. We get the following coupled equations:

$$\begin{cases} \dot{f}_{AT}(t) & = (1 - \lambda(t) - \nu\frac{Q-1}{Q})f_{AT}(t) + \frac{\nu}{Q}f_{AM}(t) \\ \dot{f}_{AM}(t) & = (-\lambda(t) - \frac{\nu}{Q})f_{AM}(t) + \nu\frac{Q-1}{Q}f_{AT}(t) \end{cases} \tag{S118}$$

where $f_{AT}(t) + f_{AM}(t) = 1$. We obtain the asymptotic fitness $\lambda(t) \to \lambda$ by solving Eq. (S118) when time derivatives are 0 (and the condition that frequencies sum to 1). We obtain an asymptotic fitness (that we can approximate in the limit $\nu \ll 1$):

$$\lambda = \frac{1 - \nu + \sqrt{(1-\nu)^2 + \frac{4\nu}{Q}}}{2} \sim 1 - \frac{Q-1}{Q}\nu \tag{S119}$$

Then, to estimate the fraction of sequences which are not $A$ at the **TR**, we consider that the main contributor corresponds to sequences from the $AT$ and $AM$ populations which mutated to the **TR**=T population (which happens at rate $\mu/Q$). This hypothesis is well justified from the exact derivation and simulations of Fig. S13. Because the rate $\nu$ is much larger than $\mu$, we make the approximation that the **VR** is immediatly replaced by a $T$ as well. This results in the following dynamics for the **TR**=T, **VR**=T population fraction, the population which lost its A in its **TR** (in this approximation):

$$\dot{f}_{TT}(t) = (1-\lambda)f_{TT}(t) + \frac{\mu}{Q} \tag{S120}$$

of solution, using Eq. (S119), given by:

$$\begin{aligned} f_{TT}(t) &= \frac{\mu}{Q(1-\lambda)}\left(e^{(1-\lambda)t} - 1\right) \\ &\sim \frac{\mu}{(Q-1)\nu}\left(e^{\frac{Q-1}{Q}\nu t} - 1\right) \end{aligned} \tag{S121}$$

such that the average fraction of population without an A in their **TR** is approximately given by:

$$\begin{aligned} f_{\text{not A}} &= \frac{1}{\tau}\int_0^\tau f_{TT}(t)\mathrm{d}t \\ &= \frac{\mu}{\nu(Q-1)}\left[\frac{Q}{(Q-1)\nu\tau}(e^{\frac{Q-1}{Q}\nu\tau} - 1) - 1\right] , \end{aligned} \tag{S122}$$

as given in Eq. (4) of the main text. We check in Fig. S16 the expression in three different regimes depending on the value of $\tilde{\nu} = \nu(1 - 1/Q)$, $\tilde{\mu} = \mu Q$ and $\tau$: $f_{\text{not A}} \sim \tilde{\mu}\exp(\tilde{\nu}\tau)/(\tilde{\nu}^2\tau)$ for $\nu \gg 1/\tau$, $\tilde{\mu}/\tilde{\nu}$ for $\nu \sim 1/\tau$, $\tilde{\mu}\tau$ for $\tilde{\nu} \ll 1/\tau$.

##### 4. Dicussion about VR=A selection by the environment

We discuss the case when the environment selects for Adenines in the **VR**.

As seen in the graph of Fig. S17 for transitions between populations, the dynamics of the population where **TR**=A remain unchanged when the environment selects for an Adenine. Consequently, the asymptotic fitness remains

$$\lambda \sim 1 - \frac{Q-1}{Q}\nu . \tag{S123}$$

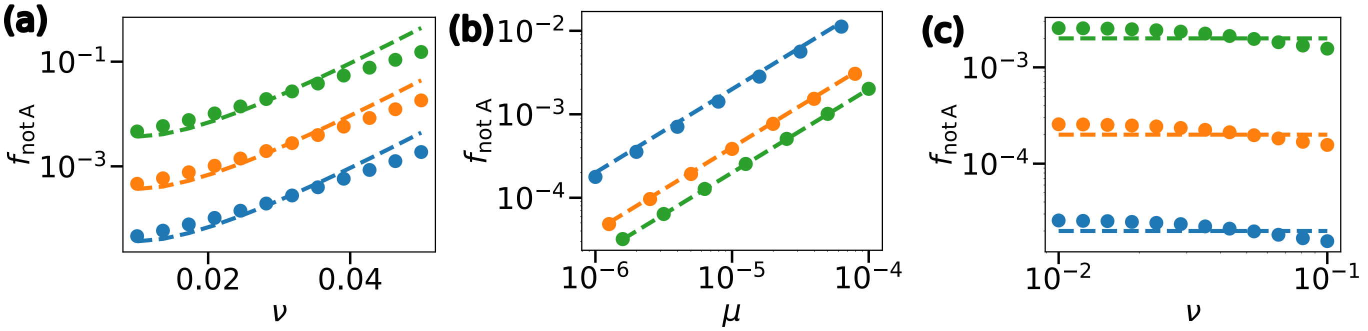


Fig. S16. Fraction $f_{not\ A}$ of sequences in the **TR** population carrying non-adenine nucleotides. **(a)** $\nu\tau \gg 1$ ($\tau = 200$, $\mu = 10^{-7}$, $10^{-6}$, and $10^{-5}$); **(b)** $\nu\tau = 1$ ($\nu = 1/\tau = 10^{-2}$, $5 \cdot 10^{-2}$ and $10^{-1}$); **(c)** $\nu\tau \ll 1$ ($\mu = 10^{-6}$, $10^{-5}$ and $10^{-4}$ and $\tau = 5$). Increasing values of $\nu$ or $\mu$ are shown in blue, orange and green successively. Dashed lines correspond to the expressions of Eq. (S122) in the respective regimes.

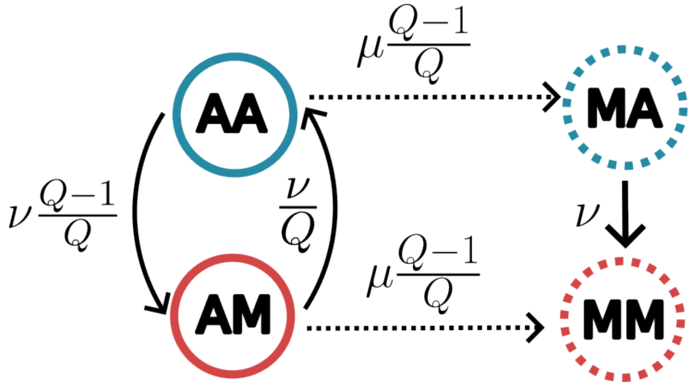


Fig. S17. **The graph of population transitions with adenine selection in the VR.** The graph of effective transitions between the different **(TR,VR)** population (first letter is the **TR**, second the **VR**). $A$ corresponds to the diversifying nucleotide, $M$ to any nucleotide except form $A$. In blue are the population with growth rate 1, in red those with growth rate 0. Hashed circles correspond to population of size of order $\mu$, while plain lines are of order 1. Dotted arrows represent transitions of order $\mu$, plain line arrows those of order $\nu$. We do not represent transitions which would lead to $\mu^2$ corrections to the population sizes.

When considering the population where **TR** $\neq A$, the dynamics of the frequencies are governed by

$$\dot{f}_{MA}(t) = (1 - \lambda - \nu)\, f_{MA}(t) + \mu\frac{Q-1}{Q} f_{AA}(t)$$
$$\dot{f}_{MM}(t) = -\lambda f_{MM}(t) + \mu\frac{Q-1}{Q} f_{AM}(t)\ . \tag{S124}$$

However, we note that both frequencies exhibit negative growth rates ($1 - \lambda - \nu \sim -\nu/Q$ and $-\lambda < 0$), preventing the expansion of either population. Thus, in this type of environmental selection, Adenines in the **TR** are not counter-selected.

## B. $L > 1$

Fot this section, we build on the intuition developed for $L = 1$: we consider that the time for the population with only $A$ at the **TR** and with the optimal sequence at the **VR** is much shorter than the time it will take to get mutations in the **TR**. Then, we consider the dynamics of the **TR** sequences which become progressively the same as the **VR** sequences. Indeed, the cost of DGR is that sequences are replaced at rate $\nu$, thus sequences with the optimal sequence directly on their **TR** have a fitness advantage of $\nu$.
We note $n_k(t)$ the number (resp. $f_k(t)$ the frequency) of the sequences with $k$ right bases on the **TR** and $L - k$ Adenines. The rest of sequences is grouped together in a single population $n(t)$ that can not make the right **VR**

sequence under DGR. Initially, $n_0(t) = N$ (resp. $f_0(t) = 1$), we only have $A$ in every **TR**. We get the following system of equation which describes the **TR** dynamics:

$$\dot{n}_0(t) = \left(L - \nu - \mu L \frac{Q-1}{Q} + \frac{\nu}{Q^L}\right) n_0 + \frac{\mu}{Q} n_1 \tag{S125}$$

$$\dot{n}_k(t) = \left(L - \nu - \mu L \frac{Q-1}{Q} + \frac{\nu}{Q^{L-k}}\right) n_k + \frac{\mu(L-k+1)}{Q} n_{k-1} + \frac{\mu(k+1)}{Q} n_{k+1} \tag{S126}$$

$$\dot{n}(t) = (L-\nu) n(t) + \sum_{k=0}^{L} \frac{\mu L (Q-2)}{Q} n_k(t) \ . \tag{S127}$$

We are interested in the regime where the population $n_0$ dominates, and define $\tau_c$ as the time where this hypothesis breaks. Thus, the equation for the frequencies become (using that the average fitness growth is mainly determined by the fitness of $n_0$):

$$\dot{f}_0(t) = \frac{\mu}{Q} f_1 \tag{S128}$$

$$\dot{f}_k(t) = \frac{\nu}{Q^{L-k}} \left(1 - \frac{1}{Q^k}\right) f_k + \frac{\mu(L-k+1)}{Q} f_{k-1} + \frac{\mu(k+1)}{Q} f_{k+1} \tag{S129}$$

$$\dot{f}(t) = \left(\mu L \frac{Q-1}{Q} - \frac{\nu}{Q^L}\right) f(t) + \sum_{k=0}^{L} \frac{\mu L (Q-2)}{Q} f_k(t) \ . \tag{S130}$$

We check in Fig. S18 that this set of differential equations describe correctly the population dynamics simulation for the population size $N$ large enough (renormalizing the frequencies to 1 at every time step to account for the dynamics after $\tau_c$ as well).

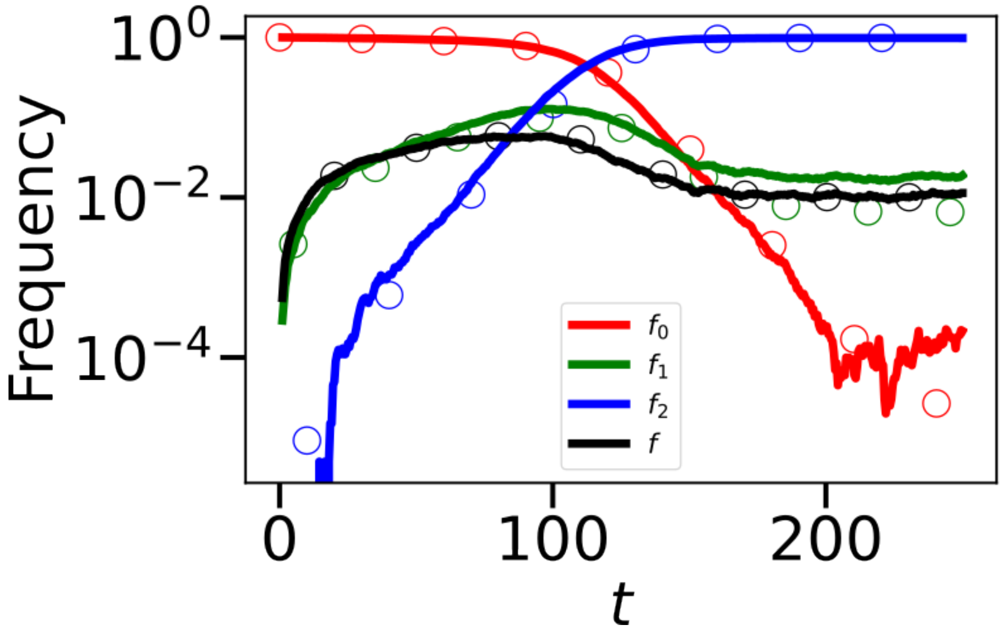


Fig. S18. **Frequency of each sub-population.** Comparison between numerical simulations (lines) and theoretical predictions (dots) of the fraction $f_k$ of the population of **TR** having $k$ correct nucleotides and $L-k$ adenines, and $f(t)$ the rest. Parameters are $L = 2$, $\mu = 10^{-3}$, $\nu = 10^{-1}$ and $N = 4 \times 10^5$.

Solving this system of equation, we get the time $\tau_c$ at which $\sum_{k=1}^{L} f_k(t) + f(t)$ gets larger than 1/2. To understand the dependance of $\tau_c$ with the parameters, we consider two situations: either sequences with higher fitness (the ones corresponding to the $f_k(t)$'s) takeover, or the length is so large that fitness advantage of adding the right nucleotides is too low and random mutations in the **TR** takeover (sequences corresponding to $f(t)$).

For the first regime, we neglect the transitions $f_{k+1} \to f_k$ which we suppose to be irrelevant to the determination of $\tau_c$. Besides,we consider $f_0(t) \approx 1$ fixed. This results in the simplified form for the triangular system of equations,

$$\dot{f}_k(t) = \frac{\nu}{Q^{L-k}}\left(1 - \frac{1}{Q^k}\right) f_k + \frac{\mu(L-k+1)}{Q} f_{k-1} \, . \tag{S131}$$

We can solve this sytem of equations in the following manner: for any $k$, $f_k(t)$ is a sum of exponentials with rates

$$\lambda_j = \frac{\nu}{Q^{L-j}}\left(1 - \frac{1}{Q^j}\right) \, , \tag{S132}$$

for $j = 0$ to $j = k$, such that

$$f_k(t) = \sum_{j=0}^{k} C_{k,j} e^{\lambda_j t} \, . \tag{S133}$$

In particular, using that for $k > 1$,

$$f_k(t) = \frac{\mu(L-k+1)}{Q} \int_0^t f_{k-1}(s) e^{\lambda_k (t-s)} \mathrm{d}s \, , \tag{S134}$$

we deduce that for $j = 0$ to $j = k-1$,

$$C_{k,j} = \frac{\mu(L-k+1)}{Q(\lambda_j - \lambda_k)} C_{k-1,j} \, , C_{k,k} = \frac{\mu(L-k+1)}{Q} \sum_{j=0}^{k-1} \frac{C_{k-1,j}}{\lambda_k - \lambda_j} \tag{S135}$$

which solution is given by

$$C_{k,j} = \frac{\prod_{m=1}^{k}\left(\frac{\mu(L-m+1)}{Q}\right)}{\prod_{m=1, m\neq j}^{k}(\lambda_j - \lambda_m)} \, . \tag{S136}$$

In particular, when only considering the dominating exponential term at every $k$,

$$f_k(t) \approx \left(\frac{\mu L}{Q\lambda_k}\right)^k e^{\lambda_k t} \, . \tag{S137}$$

Thus, $\tau_c$ is approximatively given by the smallest of the times $\tau_k$ such that $f_k(\tau_k) = 1/2$,

$$\tau_c \propto \min_{k=1,\dots,L} \left\{ \frac{kQ^{L-k}}{\nu} \ln\left(\frac{\nu Q}{\mu k Q^{L-k}}\right) \right\} \, . \tag{S138}$$

Due to the exponential dependance in $L$, we can safely assume that it is the $k = L$ term which is the minimum, leading to

$$\tau_c \approx \frac{L}{\nu} \ln\left(\frac{\nu Q}{\mu L}\right) . \tag{S139}$$

In the second regime, we consider only $f(t)$, and the $f_0(t) \approx 1$ contribution (the others are of order $\mu$ at most), which results in

$$f(t) \approx \frac{\mu L \frac{Q-2}{Q}}{\mu L \frac{Q-1}{Q} - \frac{\nu}{Q^L}} \left[ \exp\left[\left(\mu L \frac{Q-1}{Q} - \frac{\nu}{Q^L}\right) t\right] - 1 \right] \tag{S140}$$

$$\approx \frac{Q-2}{Q-2} \exp\left[\mu L \frac{Q-1}{Q} t\right] \tag{S141}$$

leading to

$$\tau_c \propto \frac{1}{\mu L} \tag{S142}$$

up to constant prefactors.
Combining Eqs. (S139) and (S142), we obtain Eq. (7) of the main text.
